# Selective decoupling of IgG1 binding to viral Fc receptors restores antibody-mediated NK cell activation against HCMV

**DOI:** 10.1101/2025.04.12.647817

**Authors:** Ahlam N. Qerqez, Katja Hoffmann, Alison G. Lee, Sumit Pareek, Kelli Hager, Akaash K. Mishra, George Delidakis, Kirsten Bentley, Lauren Kerr-Jones, Mica Cabrera, Truong Nguyen, Rebecca L. Göttler, Amjad Chowdhury, Philipp Kolb, Hartmut Hengel, George Georgiou, Jason S. McLellan, Richard Stanton, Annalee W. Nguyen, Jennifer A. Maynard

## Abstract

A key mechanism of antiviral antibodies is to bind cell-surface viral antigens and activate cellular immunity to clear infected cells, yet antibodies targeting human cytomegalovirus (HCMV) have exhibited limited efficacy. This appears due to HCMV’s multiple immune evasion mechanisms, including viral receptors (vFcγRs) which bind human IgG Fc domains to co-operatively inhibit Fc activation of host Fcγ receptors and impair Fc-mediated effector functions. We biochemically characterized and evaluated the functions of two highly conserved vFcγRs, gp34 and gp68, and mapped their binding epitopes on the Fc domain. Based on this information, we then engineered Fc variants that retain binding to CD16A, which is essential for NK activation, and to FcRn but have markedly attenuated binding to gp34 and gp68. IgG1 antibodies targeting the gB fusogen with engineered Fc domains were not internalized by infected cells, mediated enhanced CD16A activation and limited viral spread in HCMV-infected fibroblasts more effectively than wild-type Fc. Together, this work demonstrates a strategy to enhance the efficacy of antibody therapies to clear HCMV infections.

**Highlights:**

- Host and HCMV FcR compete for IgG1 binding but engage different residues.
- Fc-engineering abrogates viral FcR antagonism while retaining CD16A activation.
- Antibodies that resist vFcR capture promote superior ADCC against infected cells.
- Designer Fc domains complement Fabs to create enhanced disease-specific therapies.

## INTRODUCTION

Human cytomegalovirus (HCMV) infection is widespread and increases with age (Staras et al., 2006, Zuhair et al., 2019), with up to ~60% of the US population exhibiting seropositivity by the age of 50 (Bate et al., 2010). HCMV is usually asymptomatic in the immunocompetent but presents significant for immunocompromised individuals, including bone marrow and organ-transplant patients and neonates. In one pediatric allogeneic hematopoietic stem cell transplantation study, 49% of patients developed HCMV infection and 5% of children died from CMV complications (Sedky et al., 2014). HCMV is also the leading infectious cause of congenital birth defects (Kirby, 2016), with transmission occurring in 0.5–2% of pregnancies (Manicklal et al., 2014, Boppana et al., 2005, Kirby, 2016). Pre-emptive or prophylactic anti-viral therapies reduce clinical CMV disease in the transplant setting, but can result in myelosuppression, nephrotoxicity, and antiviral resistance (Emery et al., 2013).

The challenges of using toxic anti-viral therapeutics in patients with significant comorbidities has spurred interest in antibody-based treatments. Unfortunately, administration of intra-venous immunoglobulin containing high-titer HCMV neutralizing antibodies to pregnant women does not clearly protect the fetus (Nigro et al., 2005, Revello et al., 2014), nor is it standard treatment for transplant patients (Raanani et al., 2009). Similarly, four neutralizing antibodies targeting the fusogenic glycoprotein gB or entry glycoprotein gH (MSL109, its high affinity variant RG7667, CSJ148, NPC-21) were not effective in Phase II trials (Mokhtary et al., 2022). These efforts focused on neutralizing antibodies that prevent initial infection by blocking viral fusion with host cells and suggest that this activity alone may be insufficient to control HCMV *in vivo*. This could be because HCMV disseminates as a highly cell-associated virus (Murrell et al., 2017); accordingly, therapeutics that also target infected cells may provide greater efficacy.

In addition to neutralization, antibodies can also bind viral antigens on the infected cell surface and suppress disease by recruiting immune cells via their Fc domain to eliminate infected cells. Accumulating evidence indicating that these Fc effector functions are a crucial component of protection from HCMV. Studies using sera from gB vaccinees and seropositive women revealed strong correlations between protection and Fc-mediated antibody-dependent cellular cytotoxicity (ADCC) and phagocytosis (ADCP) (Nelson et al., 2018, Semmes et al., 2022, Semmes et al., 2023). However, these antibody activities are undermined by HCMV immune evasion strategies, mediated by viral Fcγ receptors (vFcγRs) that antagonize human Fcγ receptors – most notably, CD16A which mediates antibody-dependent cellular cytotoxicity (ADCC) by natural killer (NK) cells, as well as a wide range of other viral proteins that inhibit natural killer (NK) cell activities through other mechanisms (Weekes et al., 2014, Patel et al., 2018a).

HCMV expresses four putative vFcγRs on the surface of infected cells and in the virion envelope (Corrales-Aguilar et al., 2014b, Corrales-Aguilar et al., 2014a, Varnum et al., 2004, Atalay et al., 2002, Lilley et al., 2001, Bentley et al., 2024). Their presence is conserved in HCMV clinical isolates, with genes *RL11* and *UL118-UL119* (encoding gp34 and gp68, respectively) showing little sequence variation, whereas *RL12* and *RL13* are highly divergent (Corrales-Aguilar et al., 2014a). The two conserved vFcγRs, gp34 and gp68, bind distinct Fc epitopes and cooperate to internalize antibody/viral glycoprotein complexes, thereby clearing the infected cell surface to limit antibody-mediated effector functions (Kolb et al., 2021). Gp68 appears to have compromised a Phase II clinical trial for antibody MSL-109, which was internalization by vFcRs and incorporated into new virions (Vezzani et al., 2022, Manley et al., 2011). Conversely, removal of the three validated vFcγRs from a rhesus CMV (RhCMV) strain resulted in stronger host Fc receptor functions *in vitro* and a shorter duration of HCMV DNAemia *in vivo*. Viral clearance coincided with an increase in anti-HCMV antibodies, suggesting antibody Fc mechanisms played a larger role in the absence of vFcγRs (Otero et al., 2025b). Collectively, these results highlight the importance of Fc-mediated effector functions in viral control and the potential for vFcγRs to dampen these responses.

The conservation and complementary functions of gp34 and gp68 led us to speculate that disrupting Fc capture by these vFcγRs would restore activation of CD16A by anti-HCMV antibodies against viral cell-surface targets such as gB. Accordingly, we aimed to define the interactions between these two vFcγRs and human IgG1 and use this information to guide engineering of Fc variants that resist vFcγR capture while retaining host-FcγR interactions. Analysis of Fc variants with a range of vFcγR affinities provides insight into highly evolved CMV immune evasion mechanisms and suggests strategies to design potent anti-HCMV antibody therapeutics.

## RESULTS

### HCMV vFcγRs antagonize CD16A activation by internalizing human anti-gB antibodies

We first aimed to define the impact of the vFcγRs on CD16A activation and antibody internalization. For these studies, we used human MRC-5 fibroblasts infected with the common HCMV strain AD169 or an isogenic variant lacking expression of the three functional genes encoding vFcγRs (*RL11, RL12, UL119-118)* called Δ3; *RL13* was not considered because its expression is rapidly lost during *in vitro* culture. HMCV gB and the vFcγRs have similar expression kinetics with cell surface levels peaking 72-96 hours post-infection (Brey et al., 2018, Proff et al., 2018, Atalay et al., 2002, Weekes et al., 2014). The gB-specific antibody SM5-1 (Liu et al., 2021) binds pre- and post-fusion gB conformations on infected cells and virions but poorly activates NK cells against HCMV-infected cells (Nelson et al., 2018). The SM5-1 Fab with human C_H_1 and kappa constant domains was expressed with a wild-type human IgG1 Fc or a control mouse IgG2a Fc (SM5-1-mFc), which does not bind vFcγRs (MacCormac and Grundy, 1996). Additionally, we used Cytotect, a commercial high-titer HCMV polyclonal human immunoglobulin known to mediate ADCC as a positive control (Kolb et al., 2021, Corrales-Aguilar et al., 2014b, Vlahava et al., 2021).

We observed similar levels of SM5-1-mFc binding to MRC5 fibroblast cells infected with AD169 or Δ3 using flow cytometry and Western blot, indicating similar gB expression levels (**Fig. S1A,B**). Detection of the immediate-early antigen 1 (IE-1) and HLA expression level by Western blot further confirmed the strains’ similar behavior (MacCormac and Grundy, 1996). Infected cells were then opsonized with antibody and co-cultured with a mouse BW5147-derived reporter cell to directly monitor activation of human CD16A by mouse IL-2 secretion (Corrales-Aguilar et al., 2013). We observed significantly greater activation when Cytotect or SM5-1 was incubated with Δ3 versus AD169-infected cells (Corrales-Aguilar et al., 2013) (p<0.0001, **Fig. 1A**), with detectable signal at 72 hours post-infection (hpi) (**Fig. S1C,D)**. Overall, vFcγR expression dramatically suppresses CD16A activation by the SM5-1 antibody and supports the use of this antibody for further experiments.

**Figure 1.**
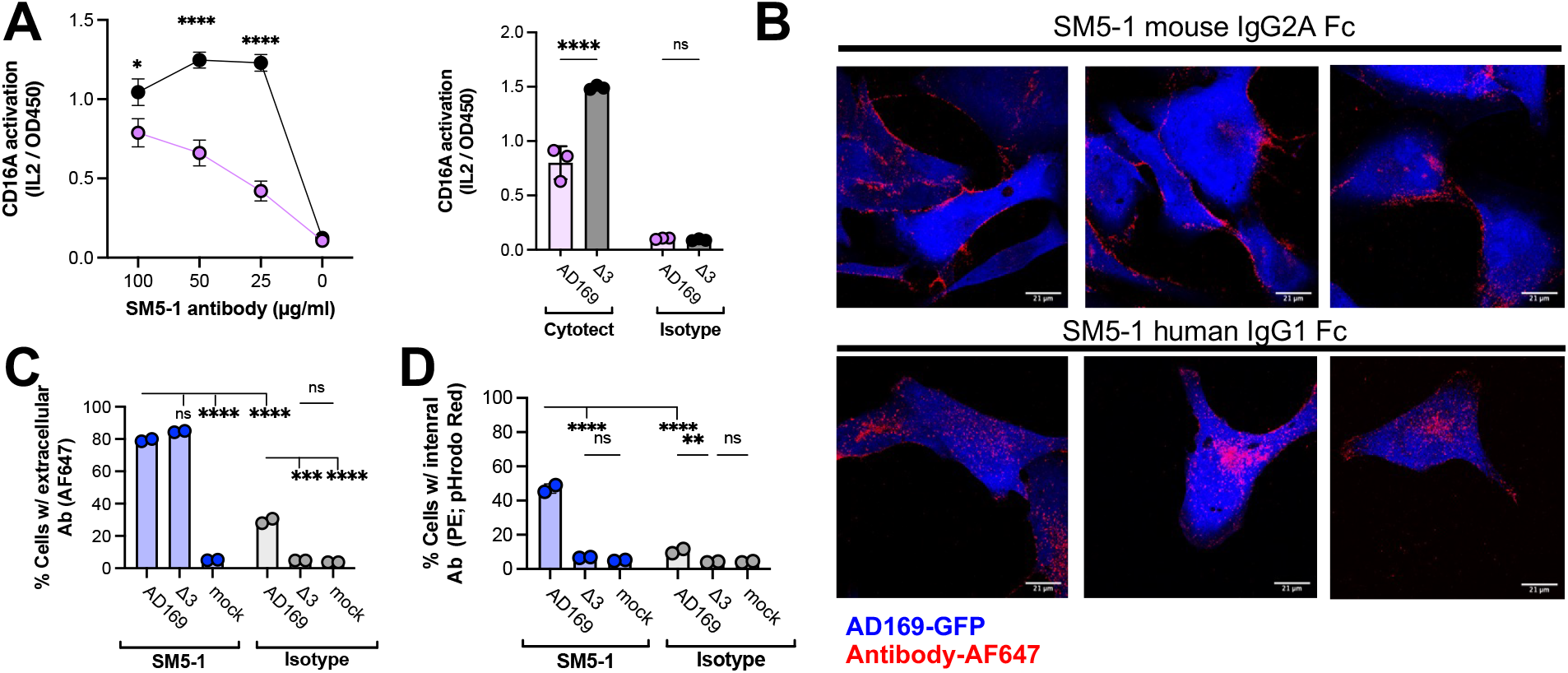
Human IgG1 antibodies are captured by HCMV-infected cells to antagonize CD16A activation. **A**, BW-CD16A-ζ reporter cells were incubated with AD169- or Δ3-infected MRC5 fibroblasts (MOI=5, 72 hpi), with CD16A activation measured by secretion of mouse IL-2 for SM5-1 and an isotype control at 100 ug/mL or Cytotect at 1:100 dilution. **B**, Fluorescent images of AD169-infected fibroblasts (MOI=2, 96 hpi) incubated with 20 μg/mL AF647-labelled SM5-1 with human IgG1 or mouse-IgG2A Fc for 2 hours at 37°C. Cells were fixed with 4% PFA and scanned using Zeiss LSM 710/Elyra S.1 at 63X. The percent of cells positive for **C**, extracellular and **D**, internalized antibody after incubation of infected cells (as above) or mock-infected cells with pHrodo-Red-labelled SM5-1 or isotype human IgG1 (67 nM) for 2 hours at 37°C was measured by flow cytometry. Extracellular antibody was detected with goat anti-human-Fcy AF647. Data are mean ± SD, for n=2 with *p<0.05, **p<0.01, ***p<0.001, ****p < 0.0001, and ns: non-significant as determined by two-way ANOVA followed by Tukey’s multiple comparisons test in GraphPad. Data are representative of one experiment; each experiment was repeated twice.

Prior work suggested that gp34 and gp68 on the surface of infected cells antagonize CD16A activation and frustrate NK cell activation by co-operating to bind human Fc domains and internalize the resulting antibody/ antigen complexes (Kolb et al., 2021). To monitor antibody binding and internalization into HCMV-infected cells, we incubated fluorescently-labelled SM5-1 and fibroblasts infected with a GFP-expressing AD169 strain before assessing antibody capture. For these studies, we selected a high antibody concentration (100 μg/ml), that is expected to be above the K_D_ for vFcγR binding. When antibodies were incubated with HCMV-infected fibroblasts at 37°C for 4 hours, confocal imaging showed that SM5-1-mFc localized to the cell periphery, consistent with the Fab region binding gB on the infected cell surface, while human SM5-1 IgG1, was primarily internalized (**Fig. 1B**). The same experiment was performed on ice to prevent endocytosis; quantification by flow cytometry showed that SM5-1 bound AD169- and Δ3-infected cells similarly (~80% cells), while an isotype control preferentially bound AD169-infected cells (~30% vs ~5% of uninfected cells; p<0.001; **Fig. 1C**), as expected for vFcγR capture of human Fc. After 37°C incubation, SM5-1 and the isotype control were internalized to a much greater extent by AD169-versus Δ3-infected cells (~50% vs 7% respectively for SM5-1, p<0.0001 for SM5-1 compared to ~10% vs 4% for the isotype, p<0.01; **Fig. 1D**). The increased binding and internalization observed for SM5-1 with AD169-infected cells supports the concept of “bipolar antibody bridging,” in which the Fab and Fc domains can simultaneously engage different HCMV viral glycoproteins and vFcγRs on the infected cell surface (Ndjamen et al., 2014). These data indicate that vFcγR expression by HCMV-infected cells mediates efficient capture of gB-specific human antibodies, which led us to hypothesize that disrupting Fc:vFcγR interactions could increase the potency of anti-gB antibody therapeutics.

### Soluble, engineered gp34 and gp68 retain Fc-binding activities

We next engineered the gp34 and gp68 ectodomains for soluble expression and used these proteins to roughly map the binding epitopes on human Fc and determine their binding kinetics. The gp68 ectodomain (residues 69–289) was cloned and produced in CHO cells as previously reported (Sprague et al., 2008), resulting in pure and homogeneous protein (t-gp68, **Fig. S2A**). The gp34 ectodomain (residues 24-182) was produced similarly but displayed high levels of aggregation on non-reducing SDS-PAGE. Under reducing conditions, gp34 ran primarily as a single band at the expected size of ~35 kDa, suggesting cysteine scrambling complicated protein production (**Fig. S2B**). Accordingly, the five cysteines in the gp34 ectodomain were individually altered to serine and screened, resulting in variant gp34-M with a C150S substitution which exhibited reduced aggregation by size exclusion chromatography (SEC) and SDS-PAGE (**Fig. S2C-D**). Notably, gp34-M measures ~35 kDa on SDS-PAGE, while SEC and static light scattering measurements indicate gp34-M forms a ~75 kDa non-covalent dimer (**Fig. S2E**). Gp34-M exhibited high levels of binding to human IgG1 by ELISA exceeding that of unmodified gp34, indicating it retains IgG1-binding activity (**Fig. S2F**). These soluble proteins appear to capture key characteristics of native gp34 and gp68 and can serve as tools to better understand Fc-vFcγR interactions.

### Human and viral FcγRs share overlapping but distinct epitopes on human IgG1

We mapped the vFcγR binding sites on Fc using soluble gp34-M and t-gp68 in ELISAs to evaluate interference with host Fc receptor binding to Fc and to evaluate binding to previously described Fc variants. These revealed that a. the gp34 epitope overlaps with that of CD16A b. the gp68 epitope overlaps with FcRn and that c. gp34 and gp68 can simultaneously bind Fc (**Fig. 2A**). vFcγR binding to Fc variants with improved FcRn affinities revealed that the YTE (M252Y, S254T, T256E) but not the LS variant (M428L/N434S) eliminated t-gp68 binding, implicating the YTE residues in the gp68 epitope (**Fig. 2B, S3A**). vFcγR binding to Fc variants that do not bind CD16A revealed that these largely retained t-gp68 and gp34-M binding, eliminating LALAPG (L234A, L235A, and P329G) and TM (L234F/L235E/P331S) (Wang et al., 2018b, Borrok et al., 2017) as potential contact residues (**Fig. 2C, S3B**). Although the host and viral receptors bind overlapping epitopes on Fc, they appear to engage at least some unique hot spot residues, suggesting that Fc variants with impaired vFcγR binding that retain host Fc receptor binding could be identified.

**Figure 2:**
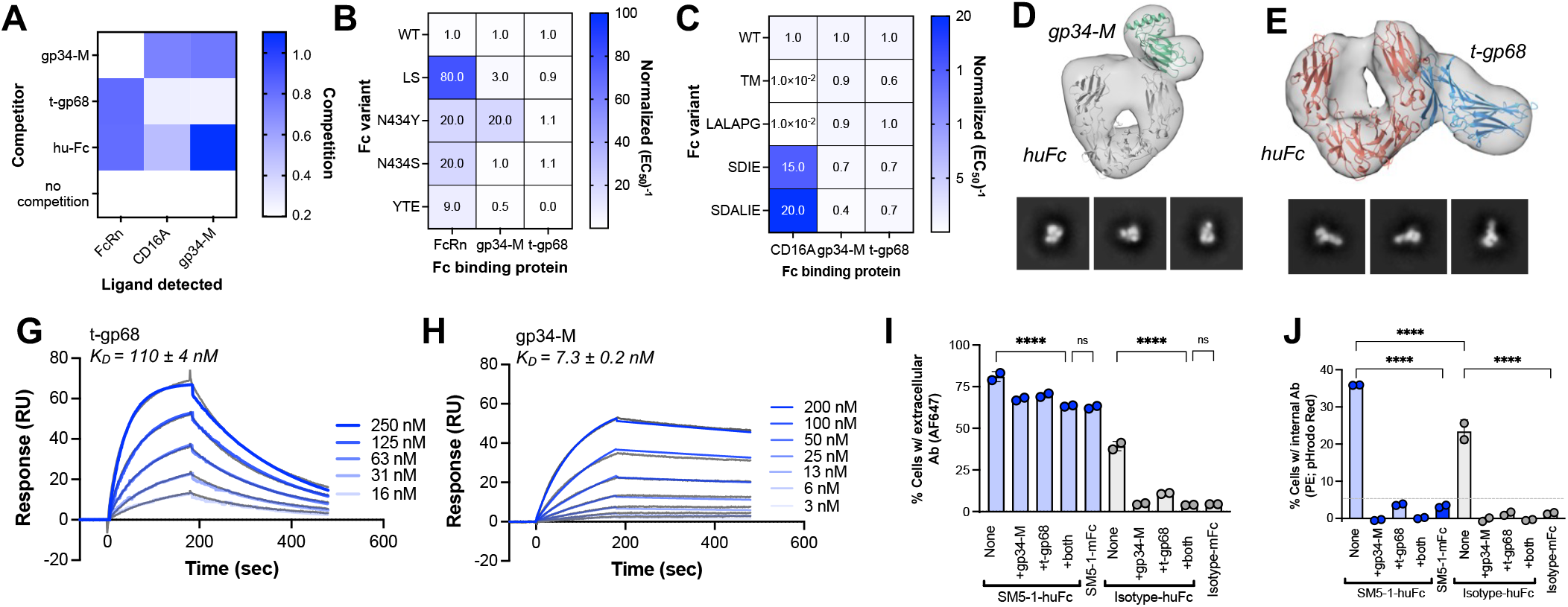
gp34 and gp68 bind human Fc with high affinity to interfere with host Fc receptors. **A**, Competition ELISA in which binding to immobilized human Fc was measured by detecting FLAG-tagged ligand (CD16A-GST, FcRn-GST, or gp34-M) in presence of a serially diluted, unlabeled competitor (gp34-M, t-gp68, hu-Fc). The assay area under the curve (AUC) was normalized to controls lacking a competitor and presented as a heat map in which darker color indicates more competition. ELISAs were used to evaluate binding of antibodies with Fc variants to gp34-M, t-gp68 and **B**, FcRn or **C**, CD16A with the normalized inverse EC_50_ values shown as a heat map such that increased color indicates increased binding. nsEM 3D reconstruction and 2D class averages for particles containing **D**, one Fc with one gp34-M bound to the CH_2_ tip and **E**, one Fc with one gp68 bound to the CH_2_-CH_3_ interface. RoseTTAFold-based gp34 (green) and gp68 (blue) models were used to fit the 3D reconstruction with the Fc crystal structure (PDB 2GJ7). Surface plasmon resonance was used to measure **F**, binding of gp68 to immobilized Fc and **G**, binding of human IgG1 antibody to immobilized gp34-M with K_D_ values determined using BIAevaluation X100 software. The ability of soluble vFcγR ectodomains (± 2 μM t-gp68 and/or 0.1 μM of gp34-M) to inhibit **J**, binding to and **I**, internalization of pHrodo-Red-labelled SM5-1 or isotype hu-Fc (67 nM) by AD169-infected MRC5 cells (MOI = 2, 96 hpi) was measured by flow cytometry. Extracellular antibody was detected with goat anti-human-Fcy AF647; in J, the grey dashed line represents threshold for detection of internalization. Data shown are mean ± SD, n=2, with *p<0.05, **p<0.01, ***p<0.001, ****p<0.0001, with ns: non-significant. All analyses performed in GraphPad using two-way ANOVA with Tukey’s multiple comparisons test. Representative data of one experiment shown; each experiment was repeated at least twice.

As a complementary approach, we used negative-stain electron microscopy (nsEM) to visualize the gp34 and gp68 binding sites on human Fc. Purified Fc, gp34-M and t-gp68 were combined in various ratios and the complexes purified by SEC before nsEM analysis (**Fig. S3C**). The 3D reconstructions and 2D particle class averages showed single and double Fc molecules with appendages (**Fig. S3D**). Analysis of particles comprising only Fc and gp34-M suggests one gp34 dimer binds one Fc on the upper C_H_2 region (**Fig. 2D**), near the CD16A binding site. In the presence of gp34-M and t-gp68, some 2D class averages showed appendages extending from huFc (**Fig. S3E**), corresponding to partially occupied t-gp68 binding sites. Analysis of these and morphologically distinct particles consisting of a single Fc molecule formed with only t-gp68 (**Fig. 2E**) suggest t-gp68 binds the C_H_2-C_H_3 interface near the FcRn binding site and that two gp68 molecules can simultaneously bind one Fc. These data provide further support for a model proposed by (Kolb et al., 2021) in which gp34 has a similar Fc-binding footprint as CD16A, while gp68 overlaps with the FcRn footprint (Sprague et al., 2008).

Finally, Fc-vFcγR binding kinetics were measured using surface plasmon resonance (**Table 1**). Immobilized Fc exposed to varying t-gp68 concentrations yielded a K_D_ value of 110±4 nM when analyzed with 1:1 stoichiometry (**Fig. 2G**), similar to previously reported values (Sprague et al., 2008) Immobilized gp34-M followed by Fc injection yielded a K_D_ value of 7.3±0.2 nM when analyzed with a 1:1 stoichiometric model, as suggested by nsEM (**Fig. 2H**). For both vFcγRs, the on-rates were modest (~5×10^4^ M^-1^s^-1^) as may be expected for proteins that co-localize on the cell surface, with intermediate (0.004±0.001 s^-1^; t-gp68) or slow off-rates (0.0004±0.0001 s^-1^; gp34-M). These binding kinetics are stronger than Fc interactions with CD16A and FcRn (~300-1000 nM depending on allotype and 500 nM at pH 5.8, respectively, (Lee et al., 2019)), indicating that vFcγRs can successfully outcompete these receptors to capture Fc.

**Table 1.**
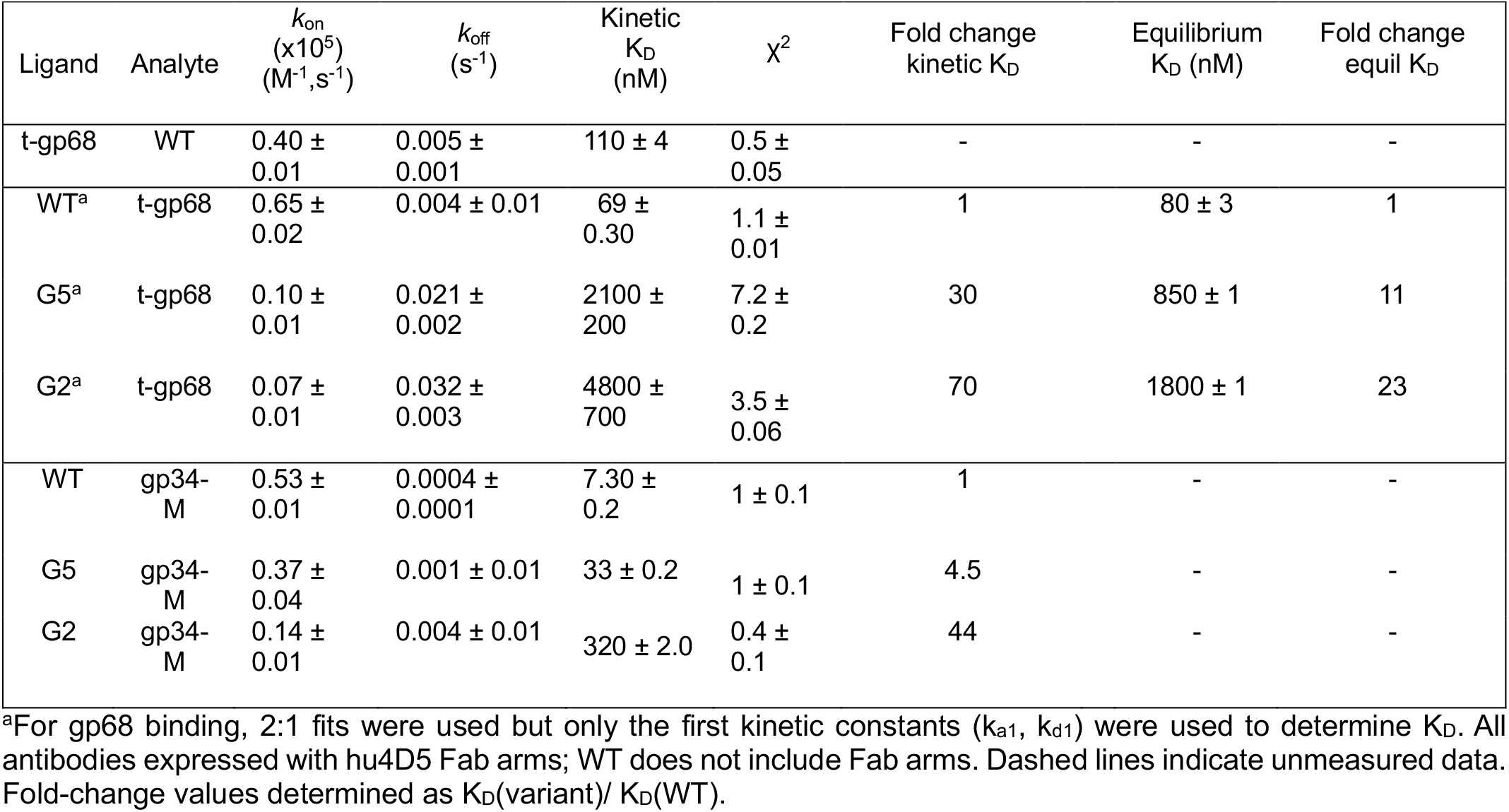
Binding affinities of Fc variants for viral FcγRs measured by SPR.

### Soluble gp34 and gp68 inhibit internalization of anti-gB antibodies by HCMV-infected cells

To further define vFcγR roles in antibody internalization, we used our soluble gp34 and gp68 ectodomains to competitively inhibit antibody capture by cells infected by HCMV strain AD169/GFP that endogenously expresses gp34 and gp68. Using our established internalization assay (**Fig. 1D**), we incubated SM5-1 or control antibodies with infected cells in the presence of soluble gp34-M or t-gp68 at concentrations of ~10×K_D_ before measuring surface-bound and internalized antibody by flow cytometry. When the incubation was performed on ice to inhibit internalization, SM5-1 interacted with gB and the vFcγRs to strongly bind AD169-infected fibroblasts (~75% positive cells), which was modestly reduced by the presence of one or both soluble vFcγRs (~60% positive cells, p<0.0001; **Fig. 2I**). When incubated at 37°C, internalization of SM5-1 and the isotype control was high (35% and 25% positive cells, respectively) but greatly diminished by the presence of soluble vFcγRs (0-5% positive cells, p<0.0001; **Fig. 2J**). Notably, while we used soluble gp34-M in this assay, gp34, RL12 and RL13 are all members of the *RL11* gene family and likely share the same binding site on Fc; accordingly, soluble gp34 may block Fc capture by all three vFcγRs. These data indicate that Fc capture is selective for antigen-bound antibodies and can be completely blocked by soluble gp34 and gp68.

### Engineered Fc domains resist vFcγR capture

We engineered human IgG1 Fc variants with greatly reduced affinities for gp34 and gp68 while maintaining binding to CD16 and FcRn. This was achieved using a competitive yeast display strategy to select for variants that retained binding to host receptors FcRn and CD16A in the presence of an excess of unlabeled gp34-M and t-gp68. The human Fc residues 221-340, spanning the hinge, C_H_2 and C_H_3 domains were fused to the c-terminus of Aga2 and presented on the yeast surface in the same orientation as on an opsonized particle (**Fig. 3A**). Yeast expressing wild-type Fc showed high binding to fluorescent CD16A tetramers by flow cytometry, which was inhibited by soluble gp34-M (**Fig. 3B**), and high binding to fluorescent FcRn tetramers at pH 6.0, which was inhibited by soluble t-gp68 (**Fig. 3C**), indicating that yeast-displayed Fc retains these key properties.

**Figure 3.**
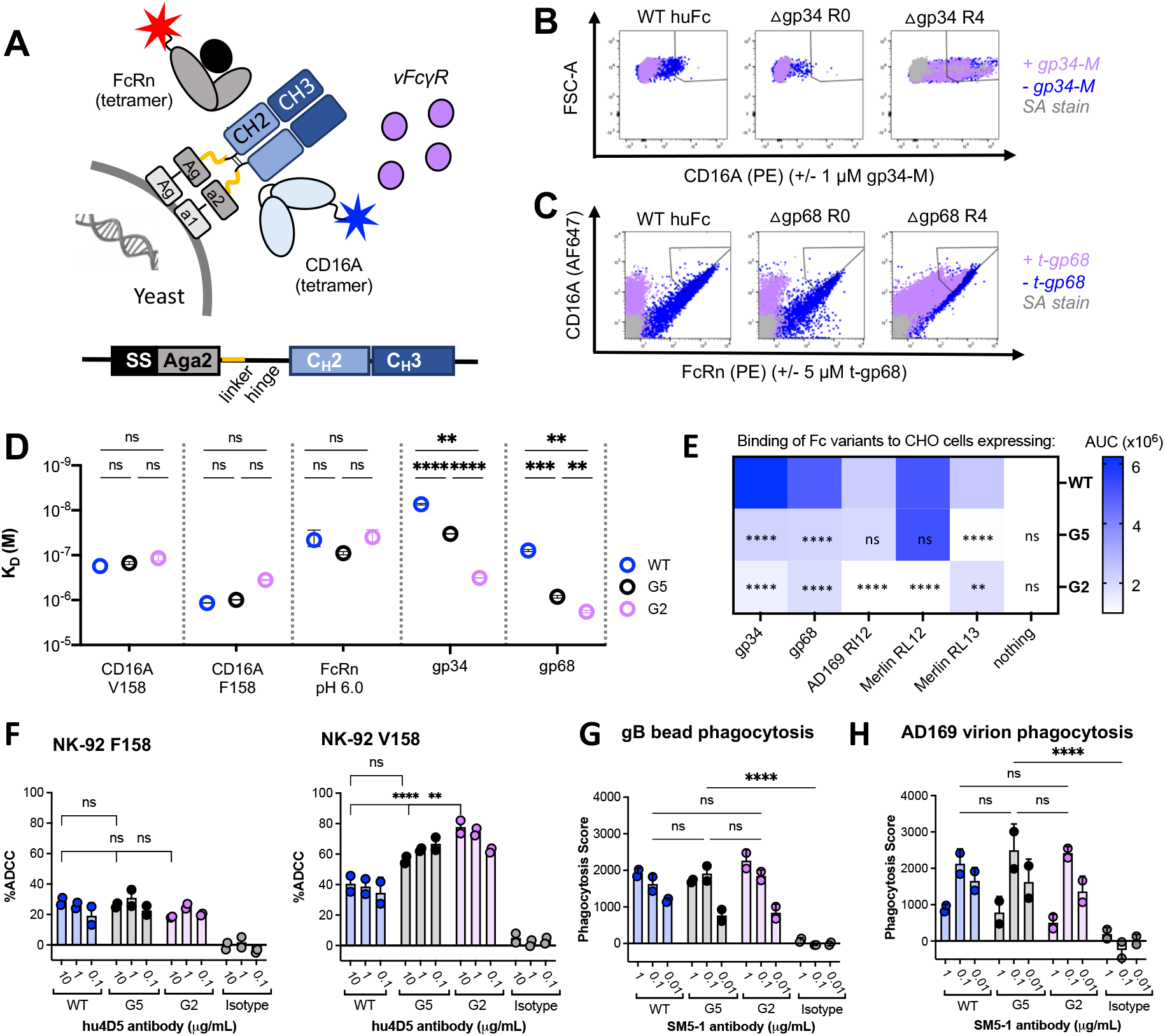
Engineered Fc domains resist vFcγR capture. **A**, Schematic of the yeast display and competitive staining strategy used to select for Fc variants. **B**, To select for gp34-resistance, an error prone CH_2_ library was displayed on yeast and clones binding soluble tetramerized AF647-CD16A (V158) in the presence of 1 μM unlabeled gp34-M were isolated by FACS. **C**, To select for clones that also exhibit gp68-resistance, a 1% error prone library was constructed from clone R47 and variants binding soluble tetramerized AF647-CD16A (V158) and PE-FcRn at pH 6.0 in the presence of 5 μM unlabeled t-gp68 were isolated by FACS. **D**, Binding affinities of Fc variants for vFcγR and host Fc receptors were determined by BLI (FcRn, CD16A V158 or F158) and SPR (gp34-M, t-gp68) with antibodies including hu4D5 Fab arms. The mean ± SEM (n=2) for K_D_ values are shown. **E**, To assess Fc binding to different vFcγRs, ectodomains were expressed on CHO cells, stained with serially-diluted hu4D5 antibodies with different Fcs (WT, G2, and G5) and detected using goat-anti-Fcγ-AF647. The mean area under the curve (AUC) is plotted as a heat map, with darker color indicating more binding. The experiment was repeated twice with n=2. For D, E significance compared to WT was assessed by one-way ANOVA with Tukey test for multiple comparisons. **F**, ADCC by hu4D5-Fc variant antibodies using HER2-positive SKOV3 target cells and NK-92 V158 or NK-92 F158 effector cells and cell lysis measured by calcein release. ADCP was performed by combining anti-gB antibody SM5-1 Fc variants with **G**, post-fusion gB-coated phrodo-Green/APC-polystyrene beads or **H**, pHrodo Green labeled AD169 virion (2 PFU per THP-1 cell) and THP1 monocytes (50 beads per cell). The phagocytosis score was calculated as the percent of APC/FITC-positive cells multiplied by the GMFI of APC and compared to WT. Representative data of one experiment shown, with each experiment repeated at least twice. Data are mean ± SD and statistical analysis was performed using two-way ANOVA followed by Tukey’s multiple comparison test, *p < 0.05, **p < 0.01, ***p < 0.001, ****p < 0.0001, ns: non-significant., **p<0.01, ***p<0.001, ****p<0.0001, ns: non-significant.

We generated a 1% error prone library of >10^6^ variants and selected cells binding CD16A-V158 tetramer by fluorescence activating cell sorting (FACS**; Fig. 3B**). This yielded two Fc variants: one (S337F) with a single S337F change and another (R47) with four changes (H268L, E294K, Q311L and K334E; **Fig. S4A**). To identify a variant with reduced t-gp68 binding and potent CD16A recruitment, we amplified residues 221-340 of variant R47 under ~1% error-prone conditions, resulting in a second library with ~10^7^ variants. Clones binding CD16A-V158 and FcRn tetramers at pH 6.0 were sorted in the presence of unlabeled t-gp68 (**Fig. 3C**). This yielded clone G2 with one additional consensus change: R255Q, which interestingly lies in between the S254T and T256E residues of the YTE changes that also reduced gp68 binding (**Fig. 2B**). We combined R255Q and S337F to generate variant G5; G5 and G2 included a Y407V cloning artifact distant from the FcγR-binding regions. Notably, the S337F and R47 changes mediating gp34-resistance and the R255Q change conferring gp68-resistance occur near the CD16A and FcRn footprints, respectively, consistent with our epitope mapping (**Fig. S4A**) and provide opportunities to evaluate the impact of Fc-resistance to a single vFcγR.

### Engineered Fc variants retain host Fc receptor interactions

These five Fc variants (G2, G5, R255Q, R47, S337F) and wild-type were expressed with different Fab arms for use in various activity assays: the control HER2-specific hu4D5 for Fc effector assays and the gB-specific neutralizing SM5-1 for anti-viral assays (Liu et al., 2021). These variants with the SM5-1 Fabs were screened for binding to viral and host FcγRs by ELISA (**Fig. S4B-D**). G2 and G5 showed the greatest reduced vFcγR binding versus WT by ELISA which was confirmed by SPR and were selected as lead clones. G5 exhibited modest changes versus WT (K_D_ increased ~5-fold for gp34; ~20-fold for gp68), while G2 exhibited dramatically reduced binding to both vFcγRs (K_D_ increased ~45-fold for gp34 and ~50-fold for t-gp68; **Table 1, Fig. 3D, S5A-B**). Additionally, G2 and G5 maintained interactions with key host FcγRs, as measured by BLI (**Table 2, Fig. S6A-C**). Binding of G2 to FcRn (at pH 6.0, no binding observed at pH 7.4) was similar, while G5 was slightly worse as compared to the values measured and in prior reports here for wild-type human Fc (Neuber et al., 2014). Conversely, G5 and especially G2 exhibited slightly increased affinity for both CD16A allotypes (**Fig. 3D**).

**Table 2.** Binding affinities of Fc variants for host FcγRs measured by BLI.

| | Equilibrium $K_D$, nM | | | |
| --- | --- | --- | --- | --- |
|  | FcRn pH 6.0 | FcRn pH 7.4 | CD16A V158 | CD16A F158 |
| **WT** | $46.0 \pm 18.3$ | ND | $174.2 \pm 8.1$ | $1160.5 \pm 14.8$ |
| **G5** | $90.3 \pm 7.6$ | ND | $150.0 \pm 11.8$ | $985.9 \pm 13.3$ |
| **G2** | $39.8 \pm 12.8$ | ND | $116.0 \pm 18.5$ | $355.8 \pm 10.5$ |

ND: not-determined kinetics due to low signal.

To assess Fc binding to viral receptors in a cellular context, we expressed the vFcγR ectodomains with *N*-terminal FLAG tags on the CHO cell surface. Expression of active gp34 and gp68 ectodomains was confirmed by anti-FLAG and human IgG1 binding using flow cytometry (**Fig. S7A**). As expected, G2 and G5 showed reduced binding to gp68 and gp34 compared to wild-type Fc (p<0.0001; **Fig. 3E, S7A**). Since the poorly conserved gp95 and gpRL13 vFcγRs are members of the *RL11* gene family with gp34, we speculated that gp34-resistant Fc variants may also lose binding to these proteins. Conserved homologies among these receptors (Cortese et al., 2012) allowed for identification of the putative immunoglobulin-fold domains for expression on CHO cells (**Fig. S7B**). G2 exhibited reduced binding to gp95 alleles from the AD169 and Merlin strains (p<0.0001) while G5 Fc had reduced binding to gpRL13 (p<0.0001) compared to wild-type human Fc (**Fig. 3E, S7A**). Fc binding to the Merlin gp95 and gpRL13 alleles was blocked by soluble gp34, suggesting these *RL11* gene family members share overlapping antibody binding epitopes (**Fig. S7C**). Lastly, we evaluated whether our Fc variants also resist capture by the HSV-encoded vFcγR homologs, gE/gI, when expressed on CHO cells. The WT Fc-containing antibody bound gE/gI in dose-dependent manner, G2 and G5 exhibited a clear loss of binding (**Fig. S7D**). Overall, we generated a panel of Fc domains ranging in affinity and specificity for multiple vFcγRs expressed by two herpesviruses.

### Engineered Fc variants retain key Fc-effector functions

We first evaluated antibody ADCC using hu4D5-containing antibodies, HER2-expressing SKOV3 target cells and GFP+ human NK-92 effector cells (Hsieh et al., 2017) expressing CD16A variants V158 or F158. For all Fc variants, the NK-92 effector cells expressing the low affinity F158 CD16a-allele resulted in ~20-30% target cell lysis. However, G2 and G5 showed increased target cell lysis versus wild-type Fc in the presence of NK-92 cells expressing the high affinity CD16A-V158 allele (60-80% versus ~30-40% respectively, p<0.001; **Fig. 3F**). Phagocytosis has also been associated with protection from HCMV. Accordingly, ADCP was evaluated using human THP-1 monocytes (CD64+/CD32A+/CD16A−) (Fleit and Kobasiuk, 1991). All Fc variants exhibited similar phagocytosis scores for gB-coated beads or AD169 virions (**Fig. 3G-H**). This indicates the engineered Fc domains can mediate key Fc-dependent effector functions associated with HCMV protection. In summary, these engineered Fcs provide an opportunity to explore the impact of high, medium (G5) and low affinity (G2) capture by vFcγRs on anti-HCMV activities.

### vFcγR-resistant Fcs mediate potent anti-viral responses against AD169-infected cells

To determine the impact of Fc engineering on anti-HCMV activities, we compared SM5-1 anti-gB antibodies bearing different Fcs in a series of assays with AD169-infected cells. When allowed to stain infected cells on ice, levels of extracellular antibody binding were similar for all variants due to Fab/gB interactions (~70%, **Fig. 4A**). However, when incubated at 37°C, antibodies with WT Fc were efficiently internalized (~35%, similar to **Fig. 1C**), this was reduced for G5 (25%) and further reduced for G2 (18%) with no internalization of the SM5-1-mFc control (p<0.010; **Fig. 4B**).

**Figure 4.**
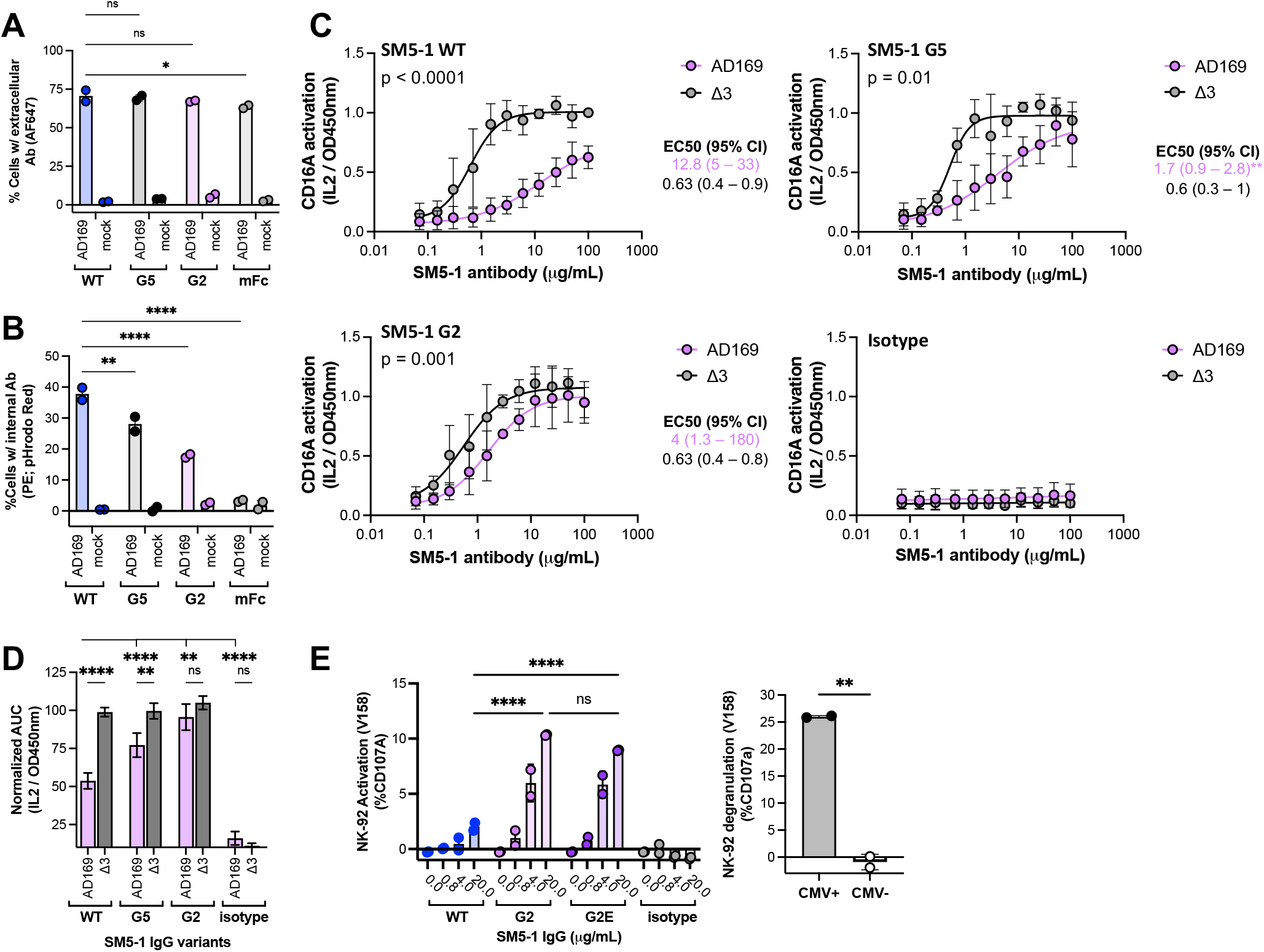
SM5-1 antibodies with engineered Fc domains mediate enhanced anti-viral activities against AD169-infected cells. **A**, Cell staining by and **B**, internalization of SM5-1 antibodies with Fc variants (20 μg/ml) in the presence of AD169- or mock-infected MRC5 cells. One-way ANOVA with Tukey’s multiple comparisons was used for statistical analysis; data are presented as mean ± SD (n=2). **C**, CD16A activation of SM5-1 antibodies measured by secretion of mouse IL-2 secretion from BW-CD16A-ζ cells in the presence of AD169- or Δ3-infected MRC5 (MOI=5, 72 hpi) and normalized to results for SM5-1 WT with Δ3. Curves from n=3 replicates were fit to 4PL or AUC using GraphPad. Mean EC_50_ for each mAb is shown and ND indicates no activation at 100 μg/mL, with p-values indicating differences in EC_50_ for activation in the presence of AD169 versus Δ3. Asterisks next to EC_50_ represent significance versus SM5-1 WT. **D**, The AUC was normalized to SM5-1 WT activation with Δ3; data shown are the mean ± SEM of 3 experiments. **E**, The percent of degranulated (CD107a-positive) NK-92 (V158) cells in the presence of AD169-infected MRC5 cells (MOI=2, 96 hpi) after incubation with SM5-1 Fc variants or polyclonal antibodies purified from CMV-negative serum or CMV+ Cytogam (30 μg/mL). Data shown as mean ± SD (n=2) and average of n=3 experiments, two-way ANOVA with Tukey’s multiple comparisons test. *p<0.05, **p<0.01, ***p<0.001, ****p<0.0001, ns = non-significant.

To determine whether IgG antibodies with vFcγR-resistant Fcs improve NK cell activities, we monitored CD16A signaling in the presence of HCMV-infected MRC5 cells using our BW-CD16A −ζ reporter cells with the high-affinity V158 CD16A allele and measured mouse IL-2 secretion. While all SM5-1 antibodies demonstrated similar levels of potent CD16A activation against Δ3-infected cells, this was greatly suppressed when SM5-1 with WT Fc was incubated with AD169-infected cells (EC_50_ values of 12.8 vs 0.63 μg/ml for Δ3 versus AD169, respectively; p<0.0001). This difference was diminished for SM5-1 with G5 (EC_50_ values of 4.0 vs 0.6 μg/ml; p<0.001) and further reduced for SM5-1 with G2 (EC_50_ values of 1.7 vs 0.6 μg/ml; p<0.01**; Fig. 4C**). The assay area under the curve (AUC) summarizes these results, showing a large activation difference against the Δ3 versus AD169 strains for the wild-type Fc which becomes insignificant for G2 (**Fig. 4D**). Analysis of additional Fc variants suggests that reduced binding to both gp34 and gp68 is required for CD16A activation, as R47 (resistant to gp34 only) or R255Q (resistant to gp68 only) were each less potent than G2 (**Fig. S8A-C**). In contrast to the enhanced ADCC observed with G2 or G5 and CD16A V158-expressing NK-92 cells **(Fig. 3D)**, no improved binding was detected in CD16A V158 binding assays (ELISAs or BLI), or immune complex reporter cell activation assays (**Fig S9A-B**), indicating the increased CD16A activation observed in response to AD169-infected cells is primarily due to reduced vFcγR capture.

During these studies, we noticed that Fc variant G2, which most strongly activates CD16A in the presence of vFcγRs, appeared to have compromised developability characteristics. This was most notable when we evaluated antibody pharmacokinetics in homozygous Tg32 transgenic mice expressing human FcRn (Avery et al., 2016) as G2 exhibited surprisingly rapid serum clearance (**Fig. S10A**). Since biochemical binding to FcRn was not impacted (**Table 2**), we evaluated antibody maternal-fetal exchange, which is also driven by FcRn-mediated transcytosis. This was performed using human trophoblast-derived BeWo cells and showed no significant differences in transport of WT versus G2 (Liu et al., 1997) (**Fig. S10B**). To determine whether we could enhance stability without altering the vFcγR-resistant phenotype, we individually reverted each change in G2 and evaluated binding to host and viral receptors. This revealed that the E294K and Y407V changes had minimal impact on vFcγR-resistance but enhanced expression (**Fig. S11A**). Accordingly, we created variant G2E retaining the H268L, R255Q, Q311L and K334E changes from G2 plus S337F to further reduce gp34 and gp68 binding. G2E exhibited similar vFcγR and host FcR affinities (**Fig. S11B**) but increased thermal stability and resistance to heat-induced aggregation relative to G2 (**Fig. S11C-D**) and restored Tg32 *in vivo* antibody pharmacokinetics (**Fig. S10A**).

To select final lead Fc variants, SM5-1 antibodies were used in an NK cell degranulation assay with NK-92 cells expressing the V158 CD16A allele and AD169-infected MRC-5 fibroblasts. The SM5-1 with wild-type Fc triggered low background levels of CD107a upregulation at high antibody concentrations (20 μg/ml), while G5 showed a slight but insignificant elevation. By contrast, G2 and G2E both mediated high degranulation rates (up to ~14%, p<0.0001) (**Fig. 4E, S11E**). Remarkably, these values for a single monoclonal antibody compare well to the degranulation observed for Cytogam (~25%), a commercial human high-titer human polyclonal IVIG preparation binding many different HCMV targets that has been evaluated clinically (**Fig. 4F**). Collectively, these experiments show an inverse relationship between reduced vFcγR-capture and enhanced CD16A activities, with G2 and G2E defining a threshold for potent CD16A activation against HCMV-infected cells.

### vFcγR-resistant Fcs mediate enhanced anti-viral activities against additional HCMV strains

To determine whether the beneficial effects of vFcγR-resistant Fcs can be generalized beyond the SM5-1 antibody and AD169 HCMV strain, we evaluated their abilities to activate CD16A with antibodies binding different gB epitopes and against other HCMV strains. For these experiments, we used SM5-1 and the non-neutralizing antibody 27-287 which binds antigenic domain 1 (AD-1) of gB (Utz et al., 1989) and stains AD169-infected HFF cells less strongly than the AD-4 binding antibody SM5-1 (**Fig. S12A**-**B**). Similar to SM5-1, 27-287 better activated CD16A signaling when combined with engineered Fc domains and AD169-infected cells than when combined with WT Fc (p<0.0001 at 50 μg/mL**; Fig. 5A, Fig. S12C**). Interestingly, activation by 27-287 with Δ3-infected cells is similar but reduced compared to SM5-1 for all Fcs, perhaps due to lower overall 27-287 binding (**Fig. S12B**). Similar to SM5-1, 27-287 mediated negligible NK cell degranulation when combined with a WT Fc and AD169-infected cells, which increased to ~8% when combined with any of the engineered Fc domains (**Fig. S12C**). This demonstrates that our engineered Fcs can enhance CD16A activation against AD169-infected fibroblasts for multiple gB antibodies.

**Figure 5.**
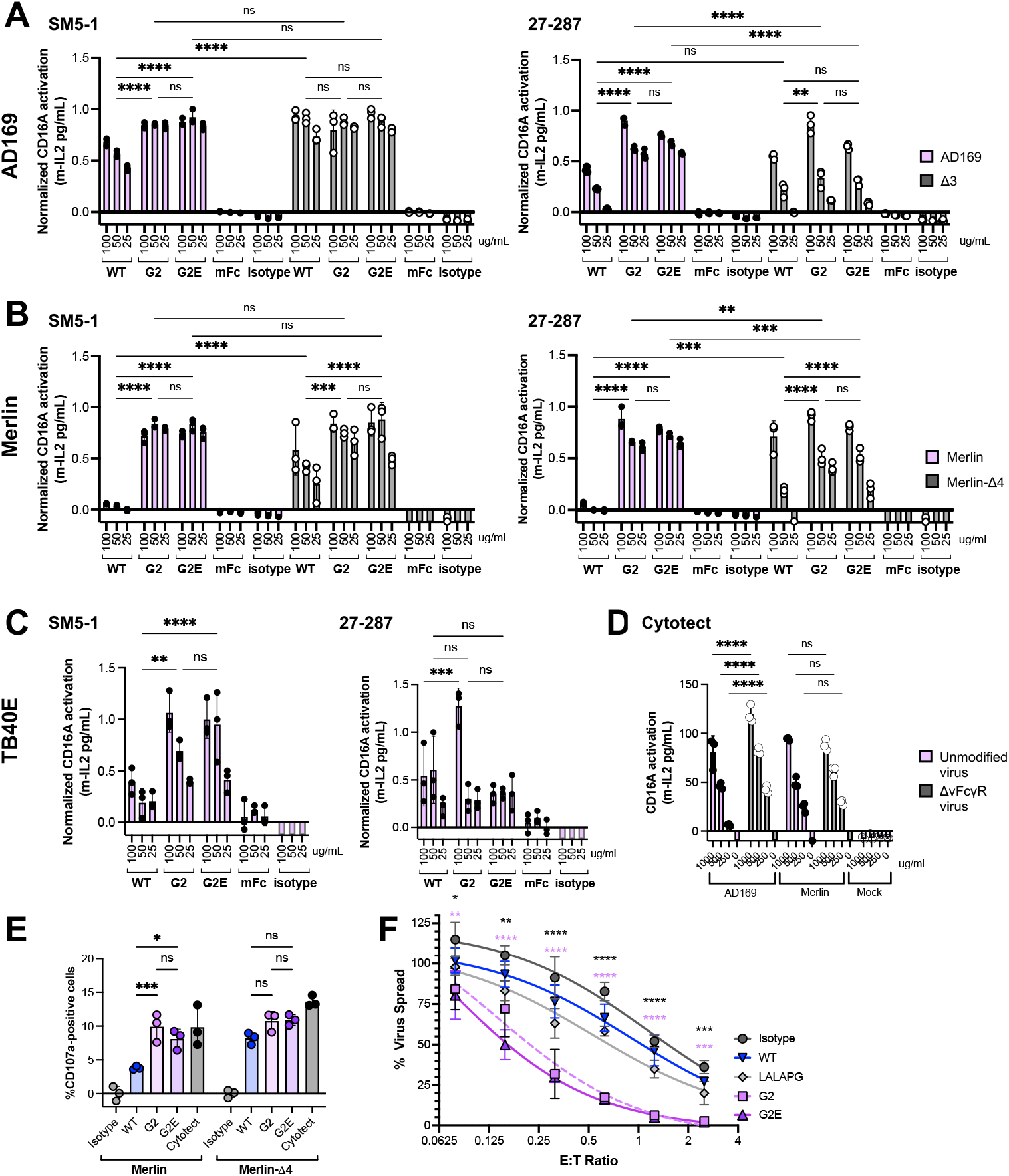
Two anti-gB antibodies with vFcγR-resistant Fcs mediate enhanced anti-viral activities against additional HCMV strains. CD16A activation by the neutralizing SM5-1 and non-neutralizing 27-287 antibodies with Fc variants was measured by secretion of mouse IL-2 from BW-CD16A-ζ reporter cells in the presence of HFF cells infected with **A**, AD169 or Δ3, **B**, Merlin or its vFcγR-deficient strain Merlin-Δ4 and **C**, TB40E, each at an MOI = 3, measured 96 hrs hpi. **D**, CD16A activation by Cytotect at 500 μg/ml with HFF cells infected by the same viral strains or a mock-infected control. Data were normalized to the maximum response per viral strain and presented as mean ± SD (n=3) of one representative experiment with one-way ANOVA with Tukey test multiple comparisons used to determine significance. **E**, *Ex vivo* purified human donor PBMC cells were combined with HFF cells infected by Merlin (pink) or Δ4 (grey) for 96 hrs (10:1 E:T ratio or 1:1 NK:T ratio) and 27-287 antibodies (20 µg/mL). The percent of NK cell degranulation (CD107a-positive) was measured after 4 hrs; data shown are mean ± SD (n=3) of one representative experiment with isotyope control subtracted from all samples. **F**, Immortalized skin fibroblasts (SFi-hTert) were infected at low MOI, with purified autologous NK cells were added at the indicated E:T ratios, along with 27-287 antibodies (5 µg/ml) at 72 hpi. Cultures were monitored in an Incucyte for 7 days, and the integrated GFP fluorescence measured for each antibody in the presence of NK cells relative to in the absence of NK cells. Each condition was assessed in technical quadruplicate and curves were fit to 4PL using GraphPad. Statistics showed compare G2E and G2 (pink and black, respectively) to WT to over different E/T ratios. Two-way ANOVA with Tukey’s multiple comparisons test was performed on all data sets (A-E, except C). *p<0.05, **p<0.01, ***p<0.001, ****p<0.0001, ns = non-significant.

We also observed enhanced CD16A activation using the clinical-like Merlin strain which expresses all four vFcγRs and the endotheliotrophic lab strain TB40E, which expresses the pentameric complex consisting of the viral glycoproteins gH, gL, UL128, UL130, and UL131 and three vFcγRs (gp34, gp68, RL12). When combined with Merlin-infected cells (**Fig. 5B)**, both SM5-1 and 27-287 showed strong CD16A activation with engineered Fcs, despite no activation by the WT Fc. When a Merlin strain lacking all four vFcγRs (Merlin-Δ4) was used, both antibodies with WT showed CD16A activation, which was enhanced by the engineered Fcs (p<0.001 for SM5-1 and p<0.0001 for 27-287 at 50 μg/mL). When combined with TB40E-infected HFF cells (**Fig. 5C**), SM5-1 with G2 or G2E enhanced activation versus WT although this was less dramatic compared with the levels seen with AD169-infected cells (**Fig. 5A**), and further diminished for 27-287 antibodies (**Fig. 5C)**. Interestingly, Cytotect showed enhanced CD16A activation in response to Δ3 versus AD169 (p<0.0001) but no difference between Merlin-Δ4and its parental strain (**Fig. 5D**). Finally, we observed enhanced antibody activity with primary human-derived NK cells against HFF fibroblasts infected with Merlin and our SM5-1 Fc variants relative to wild-type Fc at 20 μg/mL which was similar to Cytotect (**Fig. 5E**). Across three HCMV strains, our engineered Fcs consistently improved CD16A and NK cell activation relative to wild-type Fc, although the relative increase appears to vary with gB binding and altered vFcγR expression levels across strains.

To assess whether our engineered antibodies could better control cell-to-cell spread of HCMV expressing the full repertoire of immunovasins, we used the non-neutralizing anti-gB antibody 27-287 in a virus dissemination assay using Merlin. This assay provides a holistic readout for long-term virus control mediated through both cytolytic and non-cytolytic mechanisms, by measuring dissemination of fluorescently tagged virus over 7-10 days following low-level infection in the presence of NK cells and antibody (Houldcroft et al., 2020, Shan et al., 2020, Vlahava et al., 2021). To ensure appropriate MHC:KIR engagement, our assays used autologous skin fibroblasts as infected targets and primary human donor NK cells as effector cells. The NK cells with an isotype control antibody slowed virus dissemination, but only at comparatively high NK cell ratios (**Fig. 5F**). In contrast, the addition of anti-gB antibodies with engineered (G2 or G2E) but not WT or a silent LALAPG Fc enhanced NK-mediated virus control at all NK cell ratios tested (p<0.001; **Fig. 5F**). The Fc modifications allowed for enhanced control of infection, even using a clinical-like virus strain expressing four vFcγR and other NK cell-antagonizing proteins.

## DISCUSSION

Development of antibody therapeutics to treat HCMV in transplant patients and congenitally-infected infants has focused on neutralizing antibodies that block virus-cell interactions by targeting the gB fusogen or gH/gL-containing complexes that bind specific entry receptors. While these antibodies can prevent an initial infection (Blanco-Lobo et al., 2016), once established, HCMV is primarily cell-associated and spreads from cell-to-cell even in the presence of neutralizing IgG (Bournazos et al., 2014, DiLillo et al., 2016, Gardner and Tortorella, 2016, Vlahava et al., 2021, Vietzen et al., 2020, Falk et al., 2018, Murrell et al., 2017). As a result, there is a growing appreciation for the contributions of NK cells to mediate clearance of infected cells, especially for cell-associated viruses like HCMV. Moreover, modified Fc domains with enhanced CD16A binding and ADCC activities increased the potency of antibodies targeting Ebola, HIV and HCMV (Rossignol et al., 2021, Gunn et al., 2021, Van den Hoecke et al., 2017, Vietzen et al., 2020). These data, coupled with reports that HCMV vFcγRs antagonize CD16A signaling, led us to develop antibody Fc domains that better activate NK cell activities against infected cells. Accordingly, we aimed to better define Fc-vFcγR interactions and engineer human IgG1 Fc variants that resist vFcγR capture to enhance CD16A activation against HCMV-infected cells.

Since we aimed to impair Fc interactions with vFcγRs but not host Fc receptors, we reasoned that a competitive binding strategy to select clones based on host receptor binding profiles in the presence of unlabeled vFcγR would allow for use of standard yeast display technologies. We used this approach due to the absence of high-resolution structural information on Fc:vFcγR complexes. Interestingly, our selection strategy identified an R255Q substitution, which falls exactly in between two YTE changes (M252Y, S254T, T256E). Whereas YTE showed reduced gp68 binding in ELISA but also impaired CD16A affinity and ADCC (Grevys et al., 2015), R255Q reduced gp68 binding without compromising ADCC. Similarly, we identified K334E and H268L changes near the Fc/CD16A interface and Q311L which was also a substitution site in the DHS extended half-life variant (Lee et al., 2019). Protein engineering strategies were also crucial in identifying gp34 and gp68 variants for soluble expression and use in our biochemical assays. In addition to being remarkably successful against HCMV, this strategy could be adapted to engineer Fc variants that resist capture by other immune evasins, such as *Staphyloccocus aureus* protein A (Chen et al., 2022).

Antibodies with engineered Fc domains exhibited strong CD16A activities against HCMV-infected fibroblasts in multiple *in vitro* assays. We selected gB as a target since it is abundant on the infected cell surface and exhibits synchronized expression kinetics with vFcγRs (Vlahava et al., 2021), suggesting that gB-binding antibodies are impacted by these receptors. When combined with engineered Fc domains, Fabs binding two different gB epitopes mediated strong CD16A signaling and NK cell degranulation against fibroblasts infected with three different vFcγR-expressing HCMV strains (AD169, Merlin and TB40E). By contrast, the same SM5-1 and 27-287 antibodies mediated negligible NK cell degranulation when combined with wild-type human IgG1 Fc, consistent with prior reports (Nelson et al., 2018). The ability of our engineered SM5-1 antibodies to provide superior CD16A triggering can be attributed to vFcγR-resistance for AD169 and Merlin, as the wild-type and engineered Fcs have similar activities when incubated with cells infected by viruses lacking vFcγR expression (**Fig. 4,5**). Neutralizing antibody SM5-1 binds an epitope present on pre- and post-fusion gB (PDB 7KDD); this abundance likely contributes to its very strong responses as CD16A signaling is triggered by Fc clustering on the target cell surface. Non-neutralizing antibody 27-287, which binds an AD1 epitope that may only be accessible in post-fusion gB (Sponholtz et al., 2024), also exhibited improved activity with engineered Fcs, although this is consistently reduced relative to SM5-1. Since gB is primarily present in its pre-fusion conformation (Si et al., 2018), recognition of this less-abundant epitope may limit 27-287 clustering and CD16A responses. Regardless, the 27-287 antibody engineered Fcs limited dissemination of the Merlin strain over several days in the presence of human donor NK cells *in vitro* (**Fig. 5**), suggesting it may also limit cell-to-cell spread *in vivo*. Overall, engineering to reduce vFcγR capture resulted in potent antibody Fc domains that can be combined with multiple Fab arms to activate NK cells and suppress HCMV infection.

These engineered Fc domains provide insights into the contributions of vFcγRs to immune evasion. HCMV is unique among herpesviruses as the only species encoding more than one vFcγR, suggesting they may perform complementary functions (Corrales-Aguilar et al., 2014b). Kolb *et al*. proposed a model in which gp34 and gp68 cooperate to antagonize CD16A signaling and capture HCMV-specific antibodies (Kolb et al., 2021). Our kinetic and internalization data are consistent with this model: an antibody Fab binds gB, which positions it for capture by adjacent gp34 and gp68 proteins and triggers antibody internalization. The 10-fold stronger affinity of gp34 than gp68 for Fc (~7 nM and 70 nM K_D_, respectively) suggests that gp34 may act first. Moreover, since gp34 directly antagonizes NK cells by blocking Fc/ CD16A interactions, CD16A activation is sensitive to small changes in Fc/gp34 affinity, as demonstrated by the S337F variant which has just 5-fold reduced gp34 affinity but mediates increased CD16A signaling and degranulation relative to the wild-type Fc. However, engineered Fcs that resist capture by both gp34 and gp68 show the additive effects expected by the co-operative model: G5 with modestly reduced binding to both vFcγRs (~5-fold increased gp34 and ~20-fold increased gp68 K_D_) showed further enhanced activity over Fcs with only a S337F or R255Q change, while G2 with the weakest vFcγR affinities (~45-fold increased gp34 and ~50-fold increased gp68 K_D_) consistently showed the greatest CD16A signaling and NK cell degranulation activities (**Table 1, Fig. 4**). Overall, the inverse trend between Fc/vFcγR affinity and CD16A activation supports the hypothesis that vFcγRs antagonize NK cells while our engineered Fc domains provide a toolset to further define their impact on other immune functions.

Our engineered Fcs were consistently more active than their WT counterparts in the presence of Merlin HCMV strains lacking vFcγR expression, suggesting that increased activity is not solely due to vFcγR resistance. Biochemical binding assays measured a slight increase in CD16A affinity (K_D_ of 200 nM for wild-type Fc and 140 nM for G2, **Table 2**), while degranulation and CD16A activation assays performed in the absence of vFcγRs observed slight increases in basal CD16A activation. Together these data suggest the Fcs may better engage CD16A and that this may contribute to Fc potency, although this effect is much less than the dramatic activity increases for engineered versus wild-type Fcs observed with HCMV-infected cells (**Fig. 4, 5**). Interestingly, Vlahava *et al* demonstrated that independently increasing Fc affinity for CD16A can enhance ADCC against HCMV-infected cells. Monoclonal antibodies binding UL16 and UL141, two viral proteins that are expressed on the infected cell surface early after infection and before vFcγR levels rise, mediated ADCC against Merlin-infected cells when the Fc domain was modified with S239D and I332E changes that increase CD16A affinity to 2 nM (Lazar et al., 2006) without impacting vFcγR capture (**Fig. 2C**). When used in *in vitro* NK cell degranulation assays, high-affinity Fc domains conferred small increases in CD107-positive NK cells for some monoclonal antibodies when used individually, but much larger increases when a five-antibody cocktail was used (Vlahava et al., 2021). We reasoned that Fc modifications to instead reduce vFcγR binding would provide a wider therapeutic window to achieve high CD16A activation towards a clinically relevant gB epitope that is impacted by vFcγRs. Future work will examine the potential for combining changes that confer vFcγR-resistance with those that enhance CD16A binding to generate antibodies with further increased potency and to explore the synergistic effects of vFcγR-resistant antibody cocktails.

Anti-gB antibodies that activate NK cells have *in vivo* therapeutic relevance. HCMV infection is associated with a dramatic expansion of NKG2C+/CD57+ and FcγR1γ-adaptive NK cells which mediate enhanced ADCC (Wu et al., 2013). A recent serological study identified a specific correlation between ADCC activity against HCMV mediated by UL16-binding antibodies and reduced risk of fetal HCMV transmission from seropositive mothers (Semmes et al., 2023). Conversely, individuals with impaired NK immunity are particularly susceptible to HCMV disease (Biron et al., 1989, Witte et al., 2000). This indicates that antibody-mediated ADCC may be an important component of a protective immune response against HCMV but identification of appropriate viral targets and translation of these observations into an effective therapeutic antibody approach has been challenging. While gB-specific antibodies in polyclonal sera can drive ADCC responses (Vietzen et al., 2020), anti-gB monoclonal antibodies mediate very weak ADCC and NK cell degranulation *in vitro* (Vlahava et al., 2021, Semmes et al., 2023, Goodwin et al., 2020), which we hypothesize is largely due to vFcγR activities. The relevance of the vFcγR proteins for *in vivo* protection was recently demonstrated by Otero *et al* who identified and deleted gp34, gp68 and RL12 homologs from a rhesus CMV strain (Otero et al., 2025b). Infection of rhesus macaques with vFcγR-null virus showed similar initial viral loads but accelerated viral clearance during primary infection that coincided with the increase in anti-HCMV antibodies. Similarly, increased anti-CMV ADCC and ADCP was observed in rabbits immunized with vFcγR proteins (gp34, gp68, and gp95) compared to gB-only immunized animals, highlighting the potential for these proteins to augment vaccine antigens and enhance Fc-mediated effector functions (Otero et al., 2025a). Besides the vFcgR, multiple other HCMV gene products can also modulate NK cell activities by secreting inhibitors (Vlachava et al., 2023) and altering cell-surface levels of ligands for activating and inhibitory receptors (Patel et al., 2018b, Rubina et al., 2023). Some of these activities restrict ADCC specifically (Wang et al., 2018a), and also reduce cell-surface levels of entry glycoproteins to limit ADCC (Bentley et al., 2024). Merlin expresses a much wider range of these additional immune-evasins than AD169, likely explaining the greater effects of circumventing vFcγR with AD169 as compared to Merlin when assessed using *ex vivo* NK cells as opposed to CD16A reporter cells. Despite this, antibody activation of CD16A can overcome some of these strategies (Forrest et al., 2020) and the resulting IFNγ secretion can suppress virion production and cell-to-cell spread (Wu et al., 2015), as observed here.

This study provides an initial proof-of-concept for an Fc engineering approach to enhance anti-HCMV antibody activity. When combined with Fabs arms targeting two different gB epitopes and incubated with HCMV-infected cells, only antibodies with engineered Fcs mediated high levels of CD16A activation and prevented viral dissemination *in vitro*. We observed similar results with antibodies binding two different gB epitopes and three different HCMV strains, but evaluation of antibodies with further antigen specificities and subclasses will be necessary to determine the generality of these results. Future evaluation of these antibodies in limiting viremia using an *in vivo* model such as the *Rhesus macaque* will be a key step in assessing their potential clinical relevance, but the requirement for cognate Fc-host Fc receptor interactions as well as Fc-vFcγR interactions, as well as the divergence of viral homologues between species (Litvin et al., 2024) complicates the use of animal models. This work highlights the complexities of Fc biology and the potential for engineering this domain to tailor antibody activities to a given disease. Fifty years after antibody capture by HCMV-infected cells was first described (Furukawa et al., 1975), these data suggest that antibodies with engineered Fc domains may address the limitations of prior HCMV therapeutic antibodies to result in improved viral clearance and merit further evaluation.

## Acknowledgements

We thank Dr. Thomas Shenk at Princeton University for AD169-GFP and Dr. Adam Oberstein at the University of Illinois for guidance on growing HCMV in vitro. We gratefully thank Prof. Dr. med. Christian Sinzger, Prof. Dr. Florian Klein, Dr. rer. nat. Matthias Zehner and Artem Ashurov for providing HCMV TB40/E wildtype strain. This work was funded by an NSF GRFP to A.N.Q., Welch F-1767 and NIH AI181328 to J.A.M., Welch F-0003-19620604 to J.S.M., Clayton Foundation to G.G., Deutsche Forschungsgemeinschaft HE2526/9-2 to H.H. and KO6815/1-1 to P.K., Wellcome Trust grant 226615/Z/22/Z and NIH AI186964 to R.S. and Welch F-1319 for A.C. We acknowledge the University of Texas College of Natural Sciences and Cancer Prevention and Research Institute of Texas award RR160023 for support of the EM facility and award RP220587 for support of the Advanced Protein Therapeutics facility (RRID SCR_023740), both at the University of Texas at Austin.

## Author contributions

A.N.Q. and J.A.M. conceptualized the study. A.N.Q., K.H., A.G.L., A.K.M., A.C., R.G., K.B., L.K.-J. performed experiments. A.N.Q., J.A.M., H.H., P.K., J.S.M, G.G., R.S. obtained funding. A.N.Q., A.G.L., A.K.M., A.C., K.H., G.D., A.W.N. contributed to analysis. J.S.M., A.K.M., G.D. and G.G. provided sequences. A.N.Q. and J.A.M. wrote the manuscript draft, and all authors contributed to editing.

## Supplemental information

Document S1 contains Figures S1–S13 and Tables S1-S2.

## STAR Methods

### RESOURCE AVAILABILITY

Lead Contacts: Further information and requests for resources and reagents should be directed to and will be fulfilled by the lead contact, Jennifer A. Maynard.

### MATERIALS AVAILABILITY

Plasmids generated in this study will be made available on request by the lead contact with a completed Materials Transfer Agreement (MTA).

### DATA AND CODE AVAILABILITY

Any additional information required to reanalyze the data reported in this paper is available from the lead contact upon request.

## EXPERIMENTAL MODEL AND SUBJECT DETAILS

### Cell lines

The following cell lines were obtained from ATCC. CHO-K1 (CCL-61), MRC-5 fibroblasts (CCL-171), HFF fibroblasts (SCRC-1041), and SKOV3 cells (HTB-77) were maintained in complete medium (DMEM, 10% FBS, and 100U/mL penicillin) at 37°C/5% CO_2_. The human monocytic line THP-1 (ATCC TIB-202) was maintained in RPMI media (10% FBS, 100U/mL penicillin) at 37°C/5% CO_2_ and NK-92 cells expressing the V158 and F158 CD16A alleles (ATCC PTA-8836 and 8837, respectively) were maintained in α-MEM media (10% FBS (Gibco), 10% horse serum (Thermo Fisher Scientific), 0.2 mM myo-inositol (Sigma, #I7508-50G), 0.1 mM β-mercaptoethanol (Sigma, #636869), 0.02 mM folic acid (Sigma, #F8758-5G), 1.5 g/L sodium bicarbonate, 1 mM non-essential amino acids (Thermo Fisher, #11-140-050), 1 mM sodium pyruvate (Gibco# 11360-039), 2 mM glutamine, supplemented with 200 rU/mL of IL-2 (Sigma, #SRP3085-50UG), at 37°C/5% CO_2_. ExpiCHO (A29133) and Expi293 cells (A41249) were purchased from Thermo Scientific and maintained in ExpiCHO expression medium and Expi293 expression medium at 37°C, 8% CO_2_ respectively. BW5147 mouse thymoma cells (kindly provided by Ofer Mandelboim, Hadassah Hospital, Jerusalem, Israel) were maintained in RPMI (10% FBS, 0.5% pen/strep (PAN Biotech #P06-07100) sodium pyruvate (1X, Gibco #11360-039) and β-mercaptoethanol (0.1 mM, Sigma #636869)) at 37°C/5% CO_2_. BeWo b30 cells (Accegen ABC-TC535S) were cultured in DMEM supplemented with 10% FBS in a 5% CO_2_ atmosphere incubator at 37°C. Human fetal foreskin fibroblasts (HFF) and local donor skin fibroblasts (SF) immortalized with human telomerase (McSharry et al., 2008), or HFFF-hTert expressing the tetracycline repressor (Stanton et al., 2010), were maintained in complete medium (DMEM, 10% FBS) at 37°C/5% CO_2_.

### Animals

Transgenic homozygous Tg32 mice expressing human FcRn (The Jackson Laboratory Cat #014565) were used to determine antibody clearance rates *in vivo*. A total of 3-7mice were used for studies with males and females. Mice were housed in the University of Texas Animal Research Center under protocols AUP-2018-00093 and AUP-2022-00229.

### Virus stocks

The following stocks were produced and used throughout the studies: BAC2-AD169/GFP (gift from Professor Thomas Shenk, Princeton University); BAC2-AD169-varL; BAC2-AD169-varL ΔRL11ΔUL118-119 ΔRL12 (Δ3). Deletion virus mutants were generated as described previously (Kolb et al., 2021) using the primers listed in Table S1. In brief, recombinant HCMV AD169 mutants were generated according to previously published procedures (Karstentischer et al., 2006, Wagner et al., 2002) using pAD169-BAC2 (MN900952.1,(Le-Trilling et al., 2020)) corresponding to AD169varL (Le et al., 2011) as the parental genome. For the construction of the HCMV deletion mutants, a PCR fragment was generated using the plasmid pSLFRTKn (Atalay et al., 2002) as the template DNA. The PCR fragment containing a kanamycin resistance gene was inserted into the parental BAC by homologous recombination in *E. coli*. The inserted cassette replaces the target sequence which was defined by flanking sequences in the primers. This cassette is flanked by *frt*-sites which can be used to remove the kanamycin resistance gene by *FLP*-mediated recombination. The removal of the cassette results in a single remaining *frt*-site. The deletion of multiple non-adjacent genes was conducted in consecutive steps. The gene *TRL11* was deleted by use of the primers KL-DeltaTRL11-Kana1 and KL-DeltaTRL11-Kana2 The gene TRL12 was deleted by use of the primers KL-DeltaTRL12-Kana1 and KL-DeltaTRL12-Kana2. The gene UL119 was deleted by use of the primers KL-DeltaUL119-Kana1 and KL-DeltaUL119-Kana2.

Merlin was derived from a BAC encoding the complete genome of strain Merlin, which matches the genome of the original clinical virus within a patient (Stanton et al., 2010). Merlin Δ4 was constructed by deleting RL11, RL12, RL13, and UL119 by en-passant modification and has been described previously (Vlahava et al., 2021). Merlin-GFP was deleted for genes UL128 and RL13 to enable it to spread more rapidly in fibroblast cultures, and contains a P2A-GFP cassette after the UL36 gene (Nightingale et al., 2018). All Merlin viruses underwent whole-genome sequencing to verify genomic integrity following recovery from the BAC (Murrell et al., 2016).

### HCMV infection

For staining and *in vitro* assay experiments, MRC-5 or HFF fibroblasts were cultured in complete medium (DMEM (Gibco), 10% Fetal Bovine Serum, 100 U/ml Penicillin) to 80% confluency in 96-well plates prior to infection with HCMV AD169-GFP, AD169-varL, AD169-Δ3 or TB40E (Sinzger et al., 1999) at the MOI noted. Media was unchanged and virus was allowed to incubate 72-96 hours prior to performance of experiments. The pp71 plasmid and AD169/GFP BAC was a gift from the Shenk Lab. Both plasmids were electroporated into 1E6 MRC-5 cells and allowed to recover to 2-3 weeks prior to subculturing AD169/GFP containing media onto larger confluent flasks of MRC-5 (15-20 flasks). For AD169-varL strains, 15-20 flasks of MRC5 were infected at an MOI of 0.1-0.02. All infected cells were incubated for 11-14 days at 37°C/5% CO_2_ prior to harvesting supernatant and concentrating virus in media using 20% sorbitol cushion. Pellets were resuspended in filter sterilized in 7% sucrose/1%BSA/PBS and aliquots were stored at −80°C. Concentration of stored virus is determined using plaque assay or limiting dilution method.

Merlin strains were grown in HFFF or HFFF-TetR as required (Stanton et al., 2010). Flasks were infected at MOI=0.03 and incubated until 100% CPE, at which point supernatant was harvested, cells removed by low-speed centrifugation, and virus pelleted by high-speed centrifugation before being resuspended in complete medium. Aliquots were stored at −80°C and titres determined by plaque assay. To infect HFFF-hTert or SF-hTert, cells were plated at 80% confluency in DMEM lacking FCS, then infected at MOI=5 for 2h, after which cells were maintained in complete media.

