## Supplemental file 1 for "Selective decoupling of IgG1 binding to viral Fc receptors restores antibody-mediated NK cell activation against HCMV"

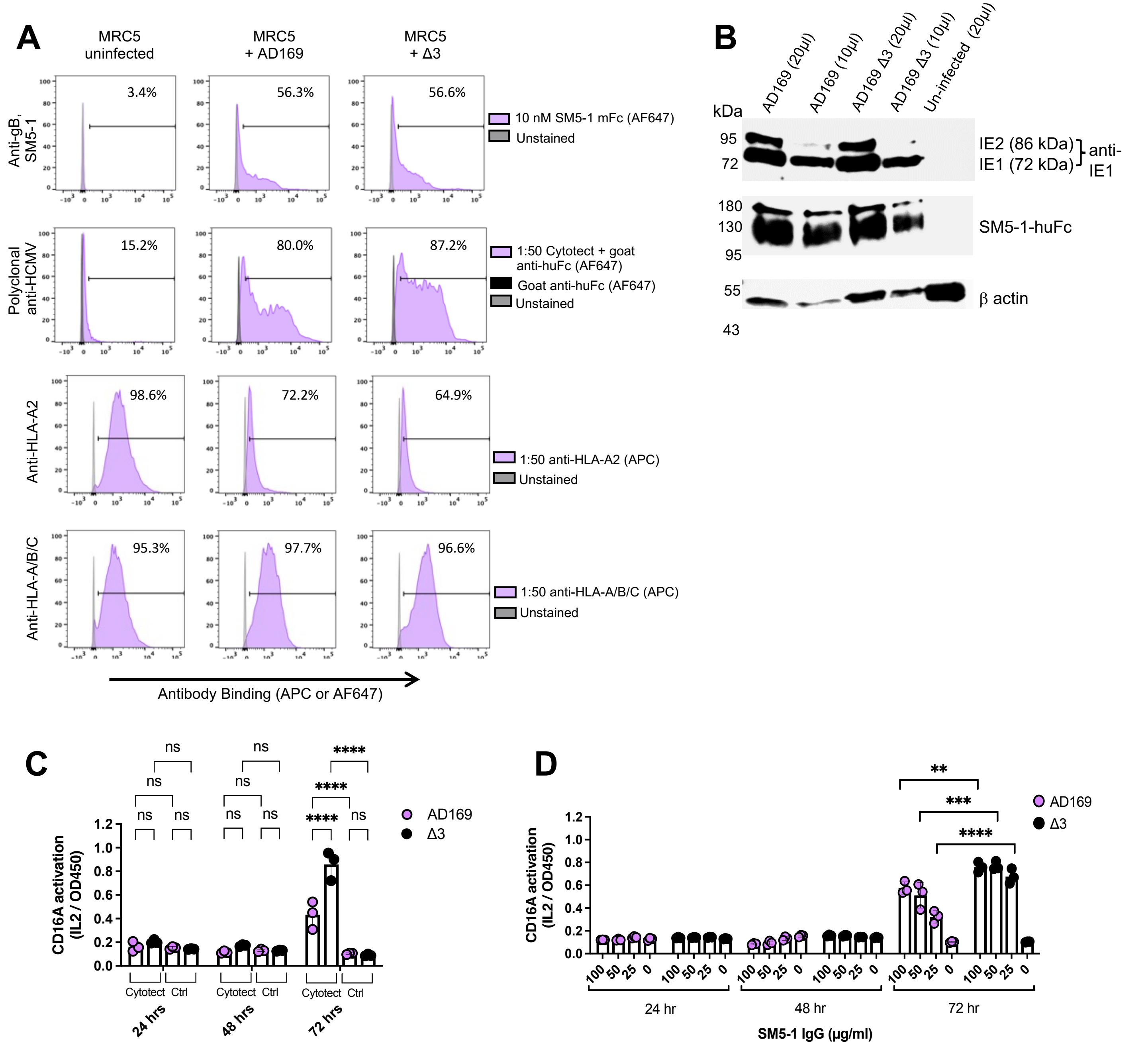


**Figure S1. AD169 and Δ3 strains have similar expression of gB and other proteins.** AD169 or Δ3 virus were used to infect human MRC5 fibroblast cells (MOI = 5, 72 hours post-infection [hpi]), while uninfected MRC5 cells were used as controls. **A,** Cells were stained with 10 nM directly labeled SM5-1-mouse Fc (AF647), 1:50 anti-HLA-A2 (APC), or 1:50 anti-HLA-A/B/C (APC) on ice or 1:50 Cytotect followed by goat-anti-human Fc-AF647. Samples were analyzed by flow cytometry, normalized histograms plotted. Positive cells were gated as indicated with the percent positive cells noted. **B,** As a complementary approach to measure gB expression, Western blot was performed on cell lysate from infected cells using SM5-1-mFc to detect gB, with IE1/IE2 expression as controls for cell infection and β actin as a loading control. CD16A activation after incubation of HCMV-infected cells with BW-CD16A-ζ reporter cells was measured at 24, 48, and 72 hpi. Secretion of mouse IL2 was measured by ELISA in the presence of **C**, Cytotect or **D,** anti-gB antibody SM5-1, each at 100 µg/mL. Data are presented as mean ± SD (n=2) repeated twice. Statistical analysis was performed by GraphPad using two-way ANOVA followed by Tukey's multiple comparison test, *$p < 0.05$, **$p < 0.01$, ***$p < 0.001$, ****$p < 0.0001$, ns: non-significant.

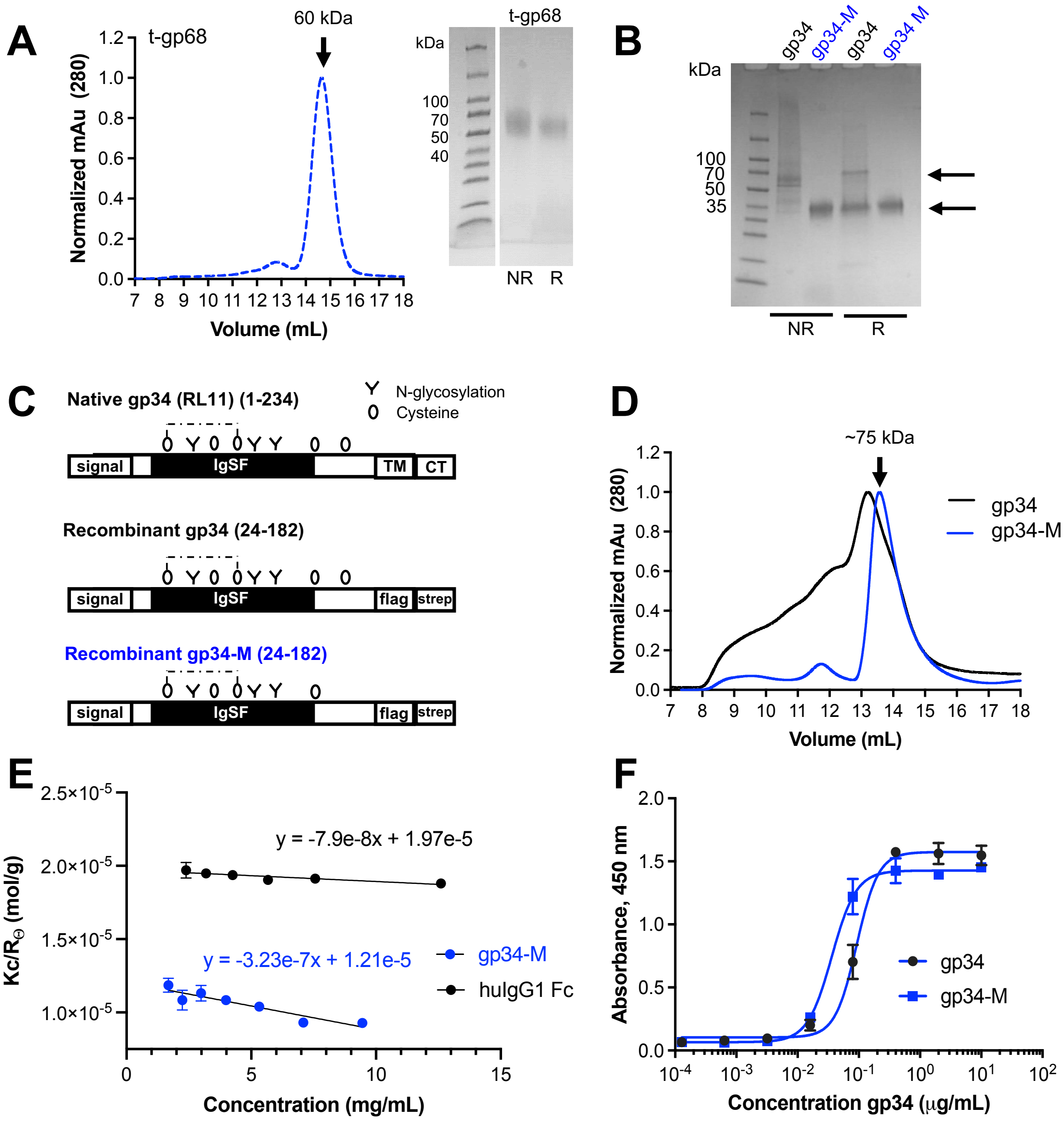


**Figure S2. Biochemical characterization of engineered gp68 and gp34 variants.** After strep tag purification and polishing with an S200 size exclusion column, 3 μg soluble ectodomain proteins were analyzed by **A,** analytical SEC and 4-20% SDS-PAGE under reducing (R) and non-reducing (NR) conditions for gp68 and **B,** SDS-PAGE for gp34 and gp34-M. **C,** Modifications of recombinant, truncated gp34 to remove N-linked glycosylation sites (Y) and cysteines (0) and generate gp34-M are shown, with the inferred di-sulfide bonding pattern (dashed line). **D,** Monodispersity of gp34 and gp34-M was assessed by injecting 100 μg purified protein on an S200 size exclusion chromatography column. **E,** Static light scattering (SLS) was used to estimate the molecular weight of purified Fc and gp34-M from the inverse y-intercept using the equation $Kc/R_\theta = 1/M_W + 2B_{22}c$; Fc was measured 51±0.3 kDa while gp34-M measured as 84±1 kDa. **F,** Fc binding activity was assessed by ELISA with immobilized human Fc, serially diluted gp34 or gp34-M, followed by anti-FLAG (M2)-HRP detection. Representative data of one experiment is shown with each experiment repeated at least twice with technical replicates.

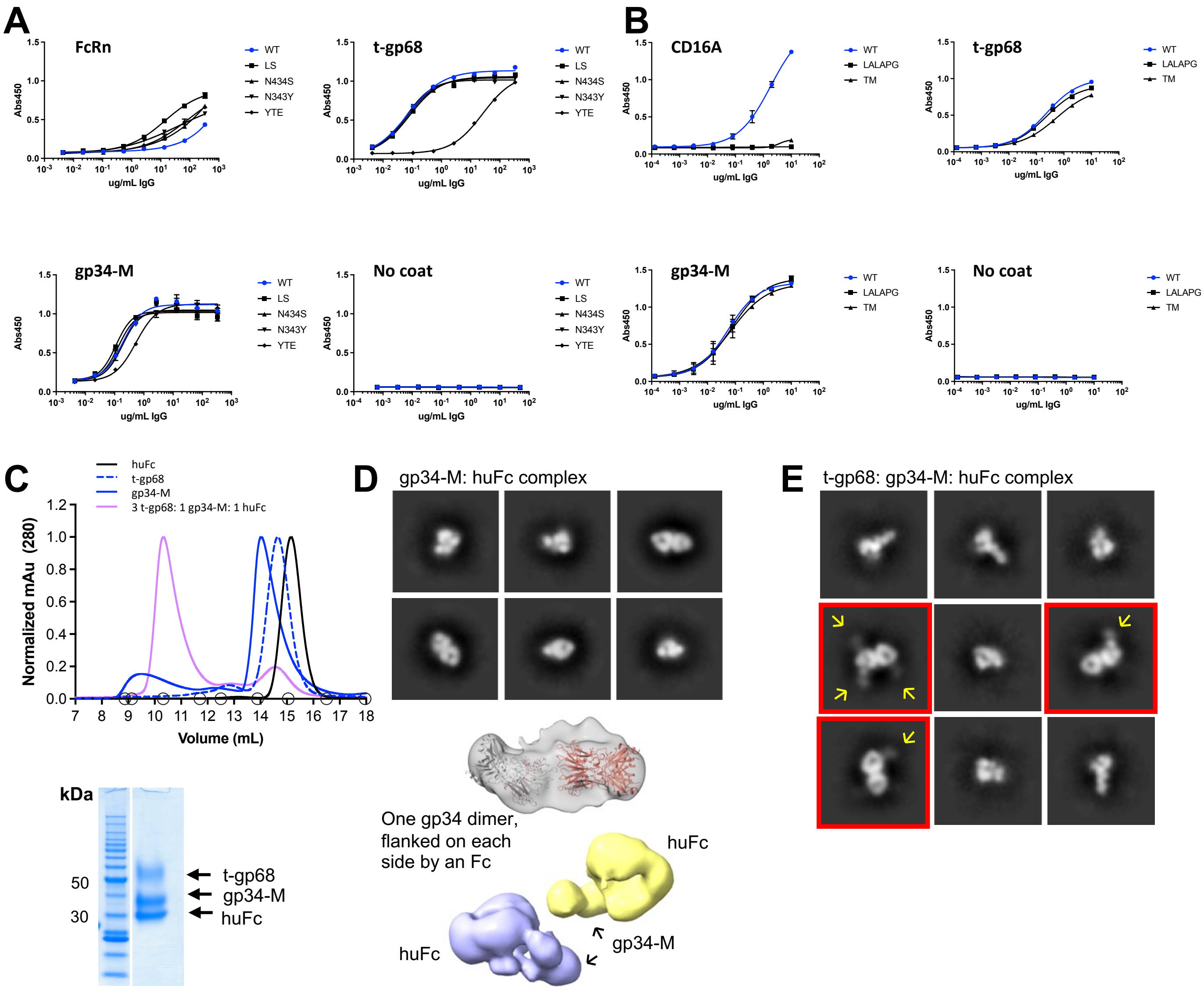


**Figure S3. Binding of Fc variants to t-gp68 and gp34-M.** ELISA with Fc variants known to have altered host receptor binding was used to determine whether these changes also impact vFcγR binding. Plates were coated with purified host Fc receptor, t-gp68, gp34-M or no coat. After blocking, serially diluted antibodies were detected with goat-anti-kappa-HRP and 4PL fits determined using GraphPad. Antibodies comprising hu4D5 Fab arms with **A,** Fc domains know to impair FcRn binding (WT, LS (M428L, N434S), N434S, N434Y, and YTE (M252Y, S254T, T256E) were assessed at concentrations from 333 to 0.004 µg/mL in 5-fold dilutions, while **B,** Fc domain with altered CD16A binding (WT, LALAPG or TM Fc domains) was assessed at concentrations from 10 to 0.0001 µg/mL in 5-fold dilutions. Representative data are shown with all experiments were performed at least twice with duplicates. **C,** Analytical size exclusion chromatography of t-gp68, gp34-M, Fc, and the ternary complex (3:1:1 molar ratio) analyzed on an S200 column and by SDS-PAGE. Molecular weight standards are indicated by circles (blue dextran, thyroglobulin, ferritin, beta amylase, aldolase, conalbumin, ovalbumin, carbonic anhydrase, cytochrome C). **D,** Negative-stain EM particle analysis of SEC-purified Fc:gp34-M complexes. *Top,* particles were used to generate negative stain 2D classes showing two Fc molecules bound together by a gp34-M dimer at the $CH_2$ tip. *Bottom,* the 3D reconstruction from the particles indicates the extra density between the two Fc molecules corresponds to a gp34-M dimer (black arrows). **E,** Negative stain EM particle analysis of SEC-purified Fc+gp34-M+t-gp68 complexes. Particles were used to generate negative stain 2D classes showing two Fc molecules bound together by a gp34-M dimer and t-gp68 appendages with partial occupancy (boxes outlined in red) protruding from the $CH_2$-$CH_3$ interface. Yellow arrows indicate partially occupied t-gp68 sites, with up to three t-gp68 molecules observed per complex.

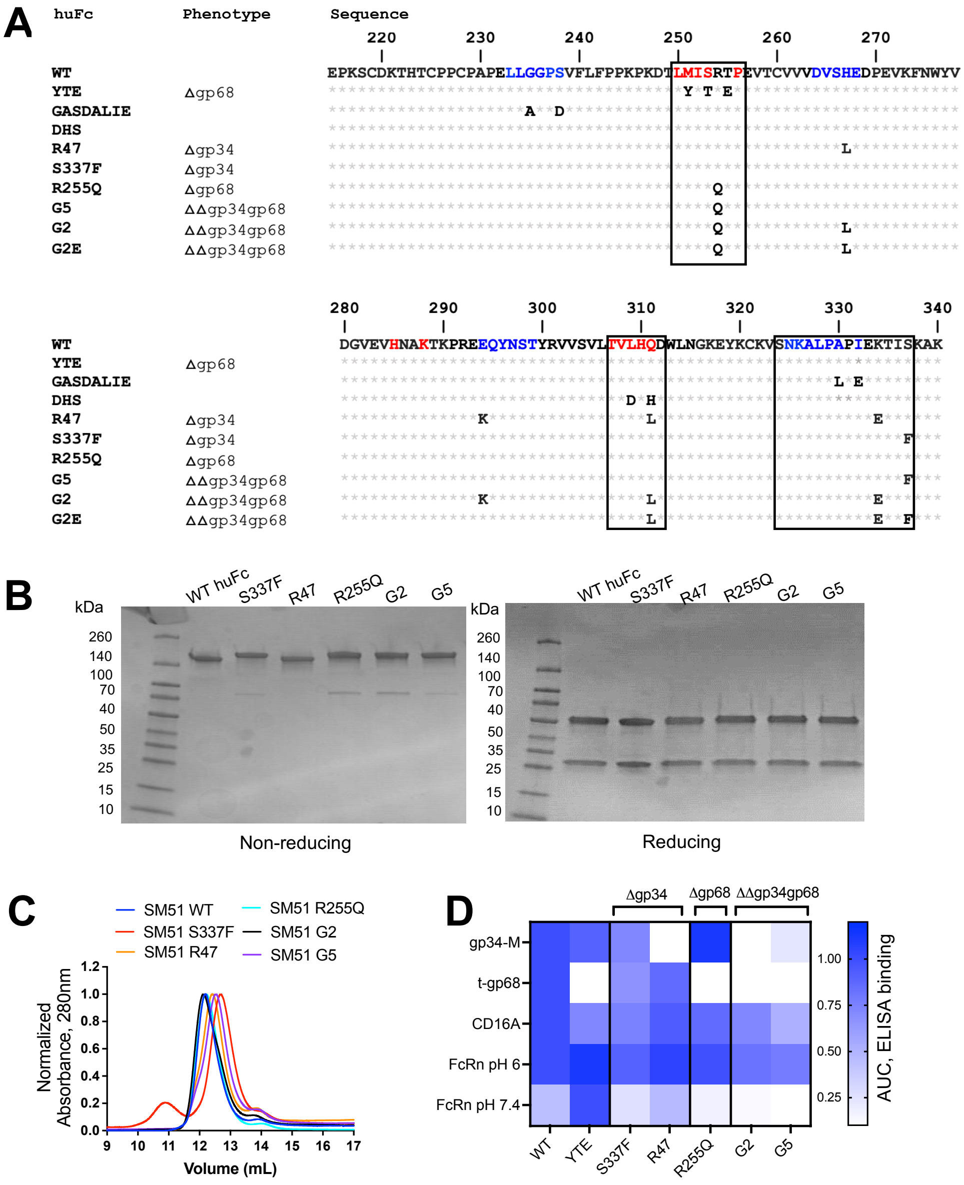


**Figure S4. Biochemical characterization of selected Fc variants. A,** Amino acid sequence allignment of $CH_2$ domains for selected Fc variants. Variants selected for reduced binding to gp34 (R47 and S337F), were combined with the variant selected for reduced binding to t-gp68 (R255Q) to generate two final clones, G2 and G5. The selected residue changes fall within the binding epitopes of FcRn (red residues) and CD16A (blue residues). After generation of SM5-1 antibodies with Fc variants, purified proteins were analyzed by **B,** 4-20% SDS-PAGE under reducing (R) and non-reducing (NR) conditions and **C,** analytical SEC to assess monodispersity after injection of 100 µg onto an S200 SEC column with normalized chromatograms shown. **D,** Antibodies with hu4D5 Fabs and Fc variants were used in ELISA to assess binding to FcRn-GST, CD16A-GST, gp34-M and t-gp68. Antibodies were coated at 4 µg/ml, followed by serially diluted Fc receptors to span the full dose-response curve and detection with anti-FLAG-M2-HRP. The area under the curve (AUC) was calculated using GraphPad, normalized to maximum AUC from the data set, and plotted as a heat map. This experiment was performed twice, representative data from one experiment are shown.

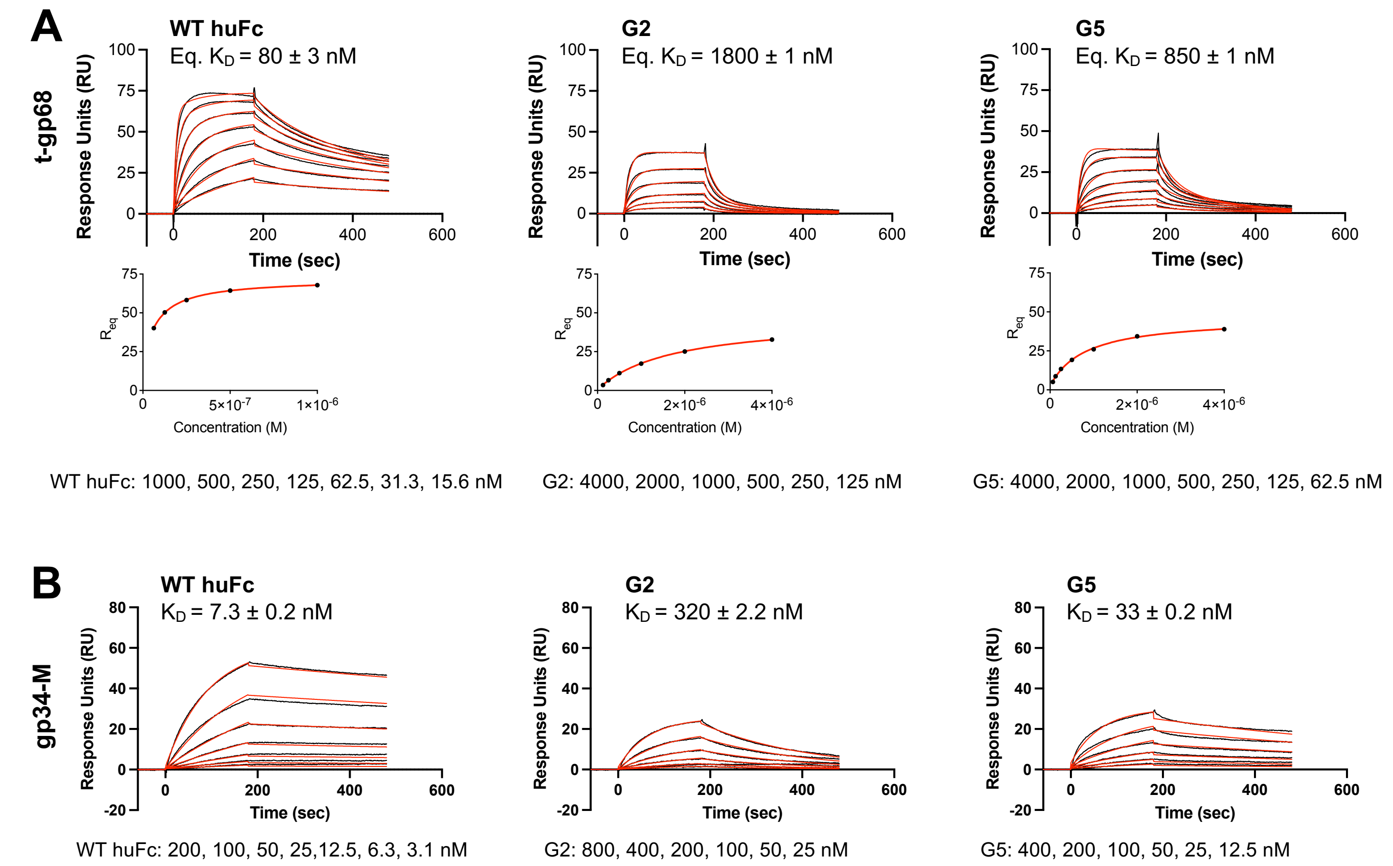


**Figure S5. Kinetics of Fc variant binding to viral Fc receptors.** SPR was performed with antibodies comprising hu4D5 Fab arms and Fc variants. CM5 chips were coupled with anti-strep Fab at 4500 RU and then t-gp68 with a twin strep tag or gp34-M with single strep tag was injected to a final RU of 35-40. Multiple concentrations (6-7) of each antibody were injected and allowed to associate for 180 seconds and then dissociate for 300 seconds. **A,** Kinetics for t-gp68 binding were evaluated with 2:1 binding fits and steady state $K_D$ values determined using the top five concentrations (1000-62.5 nM) for wild-type Fc and all concentrations (4000-62.5 nM) for antibodies with Fc variants. **B,** Kinetics for gp34-M binding were evaluated with 1:1 binding fits. All data were analyzed using the BIAcore X100 Evaluation software. Data shown are each a single representative of an experiment performed twice.

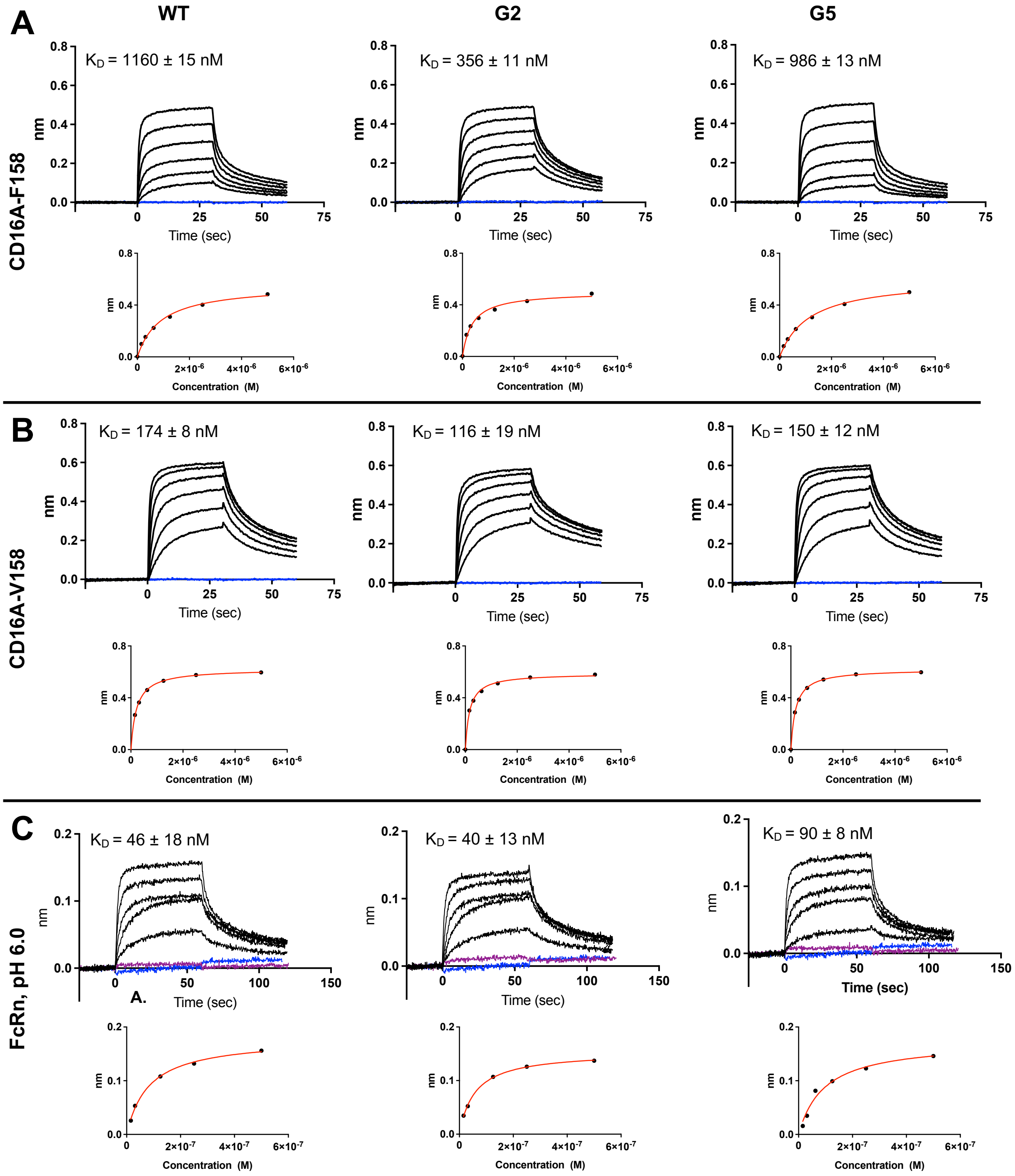


**Figure S6. Measurement of Fc binding to host Fc receptors.** BLI was performed to measure binding of hu4D5 Fc variants to the **A,** CD16A F158 and **B,** CD16A V158 alleles. For this, CH1 binding sensors were loaded with hu4D5 Fc variants (WT, G2, and G5). Loaded tips were dipped into wells containing 5000, 2500, 1250, 625, 312.5, 156.25 nM CD16A F158, or 4000, 2000, 1000, 500, 250, 125, and 62.5 nM for CD16A V158. **C.** BLI was performed to measure binding of hu4D5 Fc variants to FcRn at pH 6.0. For this, streptavidin sensors were used to capture FcRn, before dipping into wells with 500, 250, 125, 62.5, 31.25, and 15.625 nM antibody. The highest antibody concentration (500 nM) was used to assess binding at pH 7.4 (purple). In both cases, control sensorgrams lacking antibody are shown in blue. The sensograms shown are representative data from two independent runs. Equilibrium fits (red) are adjacent to sensorgrams; steady state binding kinetics were determined by Langmuir fit using Octet software and listed in Table 2.

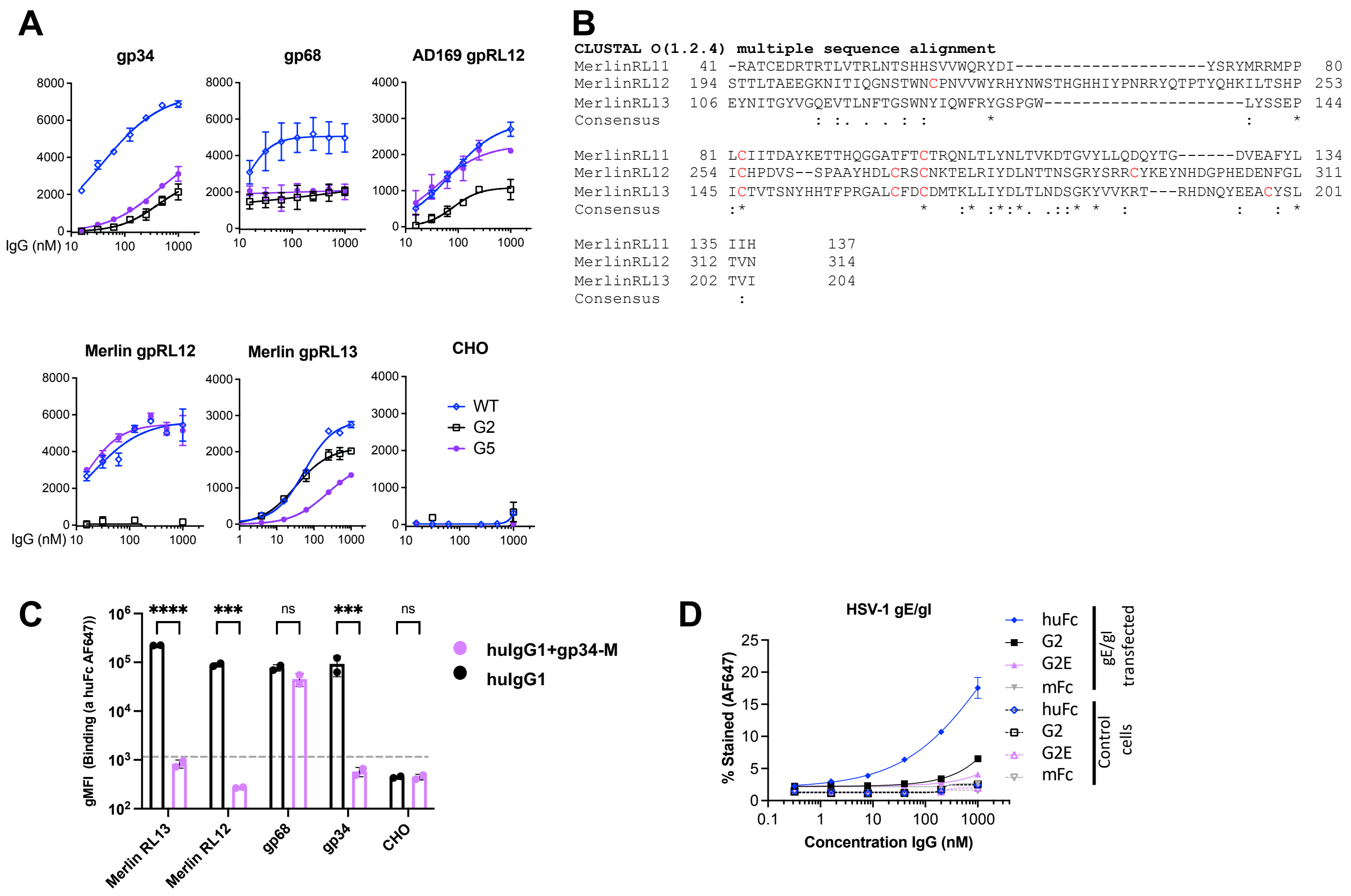


**Figure S7. Comparison of Fc binding to the four vFcγRs expressed by the Merlin strain. A,** To compare Fc binding to different vFcγRs, the extracellular domains of gp34, gp68, AD169 RL12, Merlin RL12, and Merlin RL13, were expressed on the ExpiCHO cell surface with c-terminal FLAG tags and a PDGFRα transmembrane domain. These cells were incubated with serially diluted hu4D5-Fc variants (WT, G2, and G5), then stained with anti-FLAG PE to gate on vFcγR-positive cells, and goat-anti-huFcγ-AF647 to measure the GMFI of bound antibody by flow cytometry. Double positive cells (PE+AF647) were gated and the geometric mean fluorescence intensity (gMFI) of antibody binding (AF647) plotted with 4PL curve fits performed in GraphPad. Non-transfected cells served as negative controls. **B,** Clustal Omega protein sequence alignment of the predicted Ig-fold domains of Merlin *RL11, 12,* and *13* show less than 20% homology; Ig-fold sequences were predicted using the SMART data bank. **C**, To assess the ability of gp34 to inhibit Fc binding to different vFcγRs, ExpiCHO cells expressing Merlin RL12, RL13, gp34, or gp68 were stained with 300 nM hu4D5 with WT Fc alone or in the presence of 3000 nM soluble gp34-M before measuring bound antibody as above. **D,** To assess whether the Fcs identified here also confer resistance to capture by vFcγRs expressed by HSV-1, the gE and gI ectodomains were expressed on ExpiCHOs and incubated with serially diluted hu2B1-Fc variants (WT, G2, G2E, mFc). Bound antibodies were detected by goat-F(ab)'2 anti-FcKappa-AF647 and 4PL curves fitting with GraphPad, with non-transfected cells used as specificity controls. For A, C and D, representative data of one experiment are shown, and each experiment was repeated at least twice. One-way ANOVA with Tukey test multiple comparisons used to assess significance. *p<0.05, **p<0.01, ***p<0.001, ****p<0.0001, ns: non-significant.

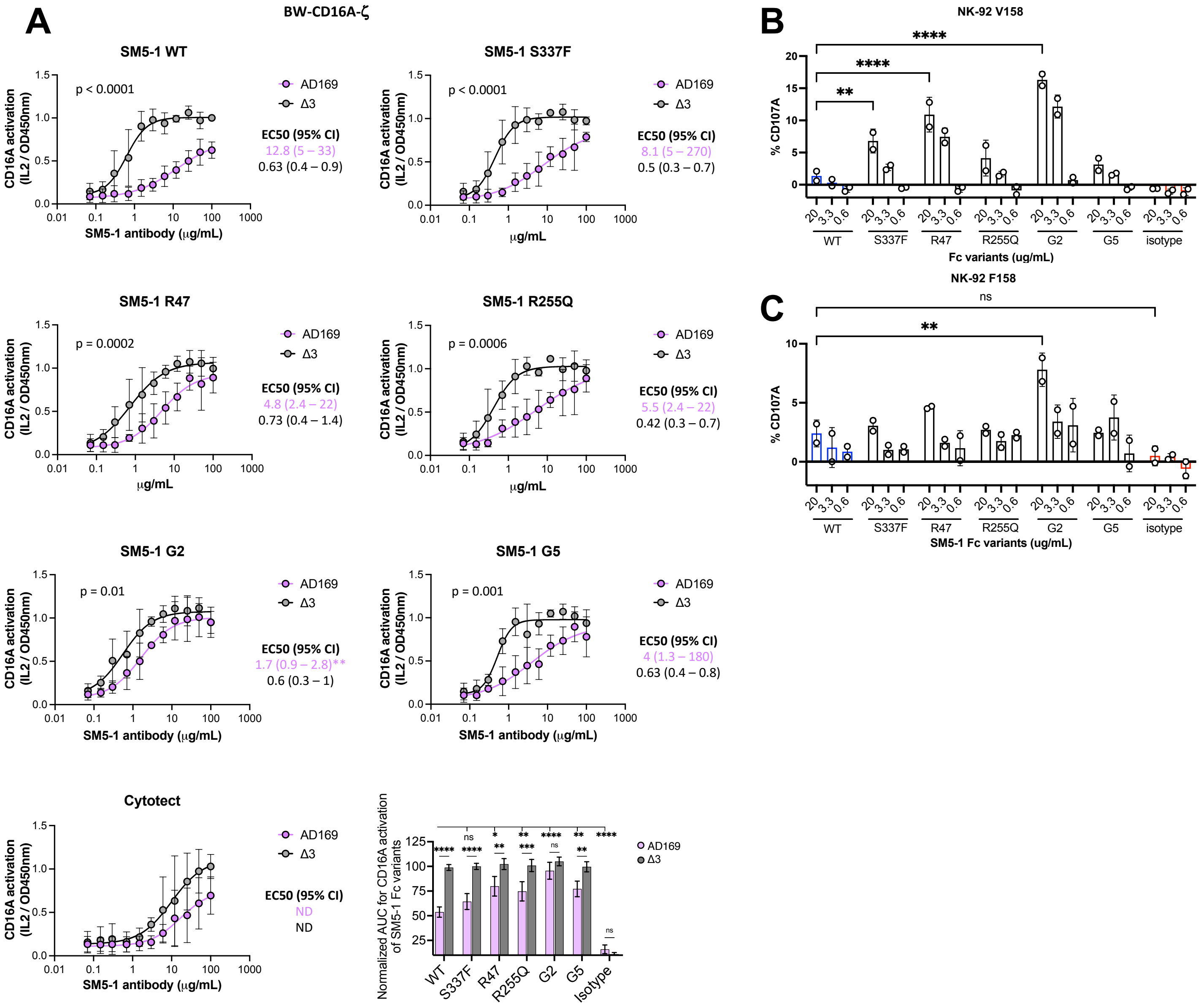


**Figure S8. CD16A activation with SM5-1 Fc variants in the presence of HCMV infected cells. A.** BW-CD16A-ζ reporter cells were incubated with AD169- or Δ3-infected fibroblasts (MOI = 5, 72 hpi) in the presence of SM5-1 Fc variants (100-0.07 ug/mL; two-fold dilutions) with CD16A signaling measured by mouse IL-2 (IL2) secretion. Data were fit to 4PL curves using GraphPad, with the mean $EC_{50}$ values (μg/mL) for each mAb or combination are noted; ND indicates activation was not detected at 100 μg/mL. The written p-values indicate $EC_{50}$ differences for the indicated antibody when incubated with AD169- versus Δ3-infected cells. Asterisks next to $EC_{50}$ values indicate the p-value for that variant versus SM5-1 WT. Also shown is the normalized AUC for each antibody normalized to that for SM5-1 WT with Δ3 infection, with analysis performed in Graphpad. Data shown are the mean ± SD of three independent experiments. The percent of degranulated NK cells in the presence of AD169-infected fibroblasts (MOI = 2, 96 hpi) and SM5-1 Fc variants (20, 3.3, 0.6 μg/mL) was measured using flow cytometry and anti-CD107A-APC for NK-92 cells expressing the CD16A **B,** V158 or **C,** F158 allele. Data presented are the mean ± SD (n=2) with statistical analysis performed using. two-way ANOVA with Tukey's multiple comparisons test Data are presented as mean ± SD (n=2) of two independent experiments and statistical analysis was performed using 2-way t-test. *p < 0.05, **p < 0.01, ***p < 0.001, ****p < 0.0001, ns = non-significant.

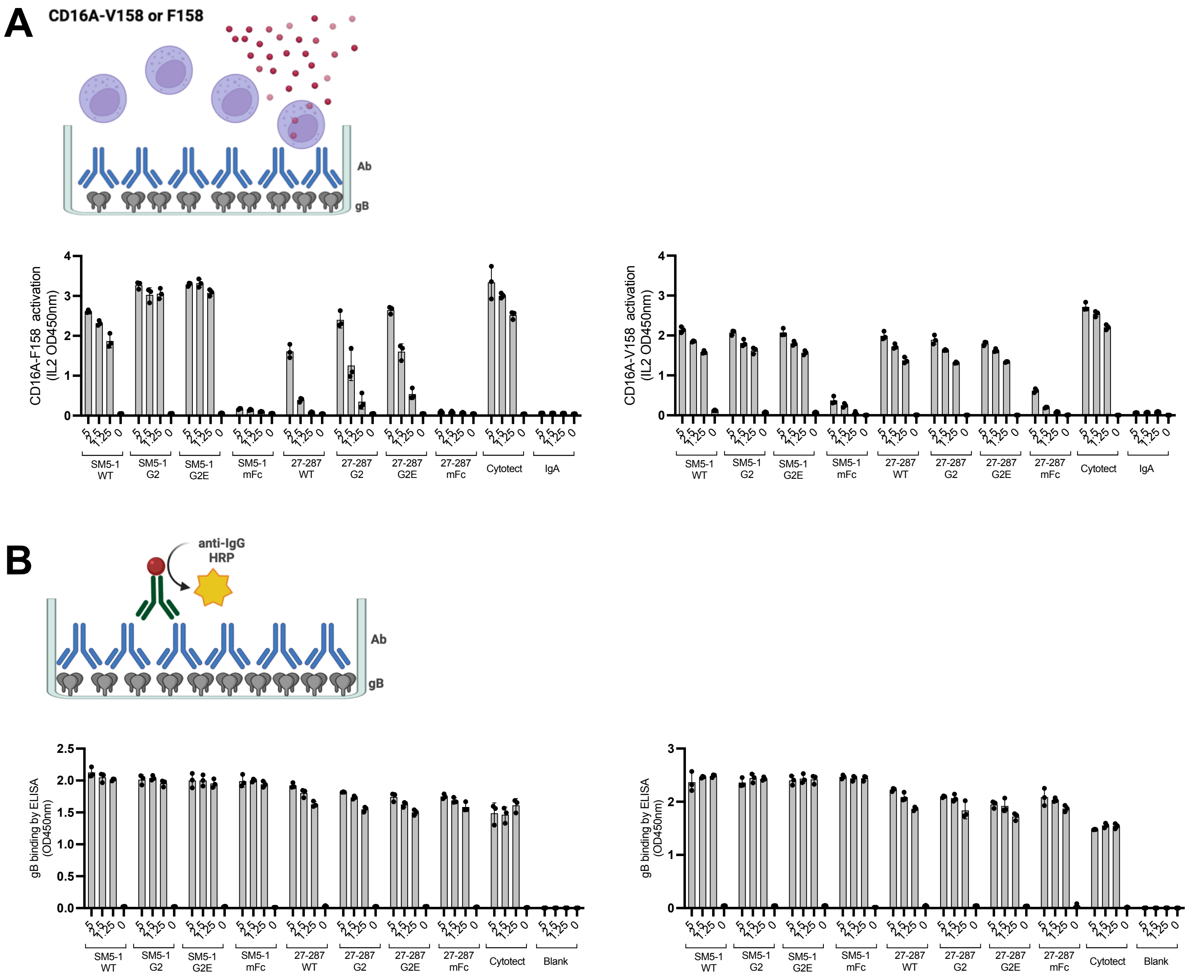


**Figure S9. Binding and CD16A activity of antibodies with immobilized recombinant gB protein.** To assess whether the antibody variants exhibit intrinsic differences in CD16A activation or gB binding, these characteristics were evaluated in a cell-free system. In both cases, purified gB protein was immobilized on Nunc MaxiSorb Elisa plates (0.5 µg/ml in 50µl PBS/well) and then serially diluted SM5-1 with different Fcs was added (two-fold dilutions starting at 5 µg/ml). After washing to remove unbound antibodies, BW-CD16A-ζ reporter cells expressing human CD16A-V158 or CD16A-F158 were added at 100,000 cells per 200µl media/well and allowed to incubate overnight. **A,** CD16A activation was measured by secreted mouse IL2 levels, using a mIL2-sandwich ELISA. Data are indicated as the average of three technical replicates from one representative experiment. **B**, Antibody bound to immobilized gB antigen was determined by adding goat-anti human IgG-HRP or goat anti-mouse IgG-HRP, respectively. Data shown depict three technical replicates from one representative experiment.

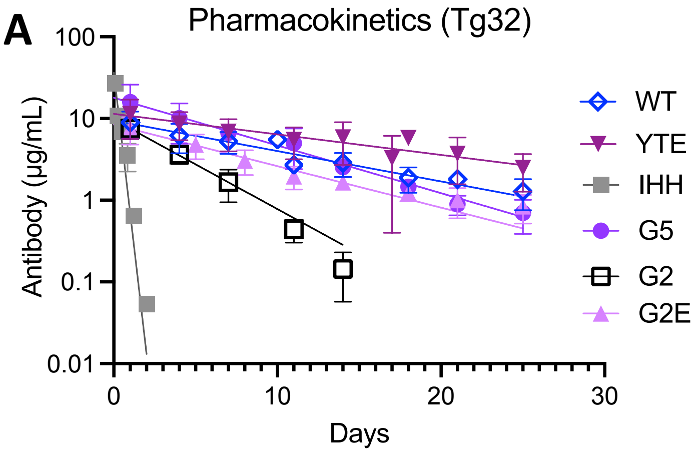


| Fc variant | $t_{1/2}$ beta (days) | p-value vs WT |
|---|---|---|
| WT | 8.8 ± 0.9 | |
| YTE | 12.0 ± 2.6 | <0.05 |
| IHH | 0.22 ± 0.01 | <0.0001 |
| G5 | 5.1 ± 1.4 | <0.05 |
| G2 | 2.2 ± 0.2 | <0.0001 |
| G2E | 7.6 ± 0.7 | ns |

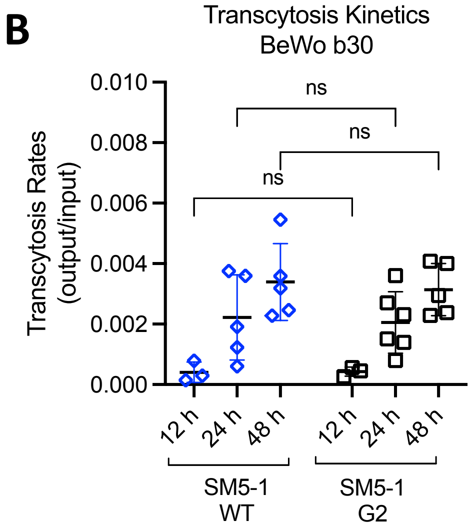


**Figure S10. Fc engineering impact on *in vivo* pharmacokinetics and FcRn-mediated transcytosis. A,** Tg32 mice with homozygous expression of human FcRn were administered 2 mg/ kg of one hu4D5 antibody with an Fc variant intra-peritoneally. The sera concentration of antibody was measured by ELISA at multiple time points with the mean ± SD shown (n=3-7). Antibody elimination half-lives were calculated by fitting data from each mouse to a single exponential decay model in Graphpad, with the rate of decay recorded as the beta elimination rate and the half-life as $t_{1/2}$ = ln2/ rate. Half-lives for each mouse administered the same Fc variant were averaged, with the mean and standard deviation tabulated. **B,** Transcytosis kinetics were performed using the BeWo b30 syncytiotrophoblast cell line, with antibody levels quantified by ELISA. Data are presented as mean ± SD (n=5). One-way ANOVA with Tukey's multiple comparisons was used for statistical analysis with * p<0.05, ** p<0.01, *** p<0.001, **** p<0.0001, and ns = non-significant.

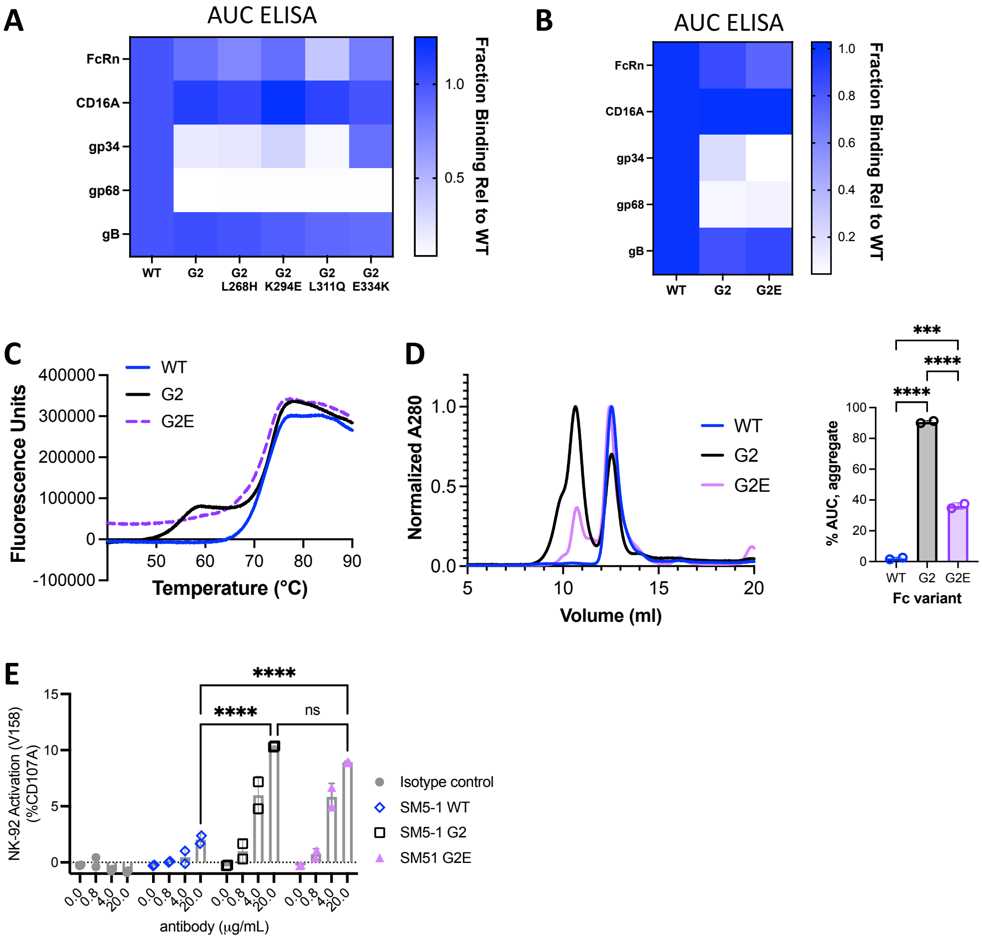


**Figure S11. Impact of individual Fc changes on receptor binding and antibody aggregation.** ELISA was used to determine whether SM5-1 G2 reversion variants retained binding profiles for gB, gp34, gp68, FcRn, and CD16A. Plates were coated with purified SM5-1 Fc variants, then serially diluted receptors (pH 7.4, pH 6.0 for FcRn) were added and detected with goat-anti-FLAG-HRP with 4PL and AUC fits determined using GraphPad. The AUC values were normalized to wild-type controls per receptor and graphed as a heat map. **B,** SM5-1 antibodies with G2 or G2E were evaluated by ELISA and graphed as function of AUC, as in A. Representative data are shown with all experiments performed at least twice with duplicates. **C,** Antibody thermal stability was assessed by differential scanning fluorimetry using SM5-1 antibodies with WT, G2, and G2E. **D,** Analytical SEC with an S200 column was used to detect the presence of higher-molecular weight aggregates after 24 hrs incubation of a 1 mg/ml at 50°C for SM5-1 antibodies with WT, G2, and G2E. The AUC of the left peak was quantified and plotted to compare aggregation. **E,** The percent of NK-92 cells (CD16A V158) that exhibited CD107A-positive degranulation in the presence of AD169-infected MRC5 (MOI=2, 96 hpi) and SM5-1 antibodies with WT, G2 and G2E are shown. Data are presented as mean ± SD (n=2) and representative of one experiment repeated twice. For D and F, one-way ANOVA with Tukey's multiple comparisons was used for statistical analysis. p<0.05, **p<0.01, ***p<0.001, ****p<0.0001, ns = non-significant.

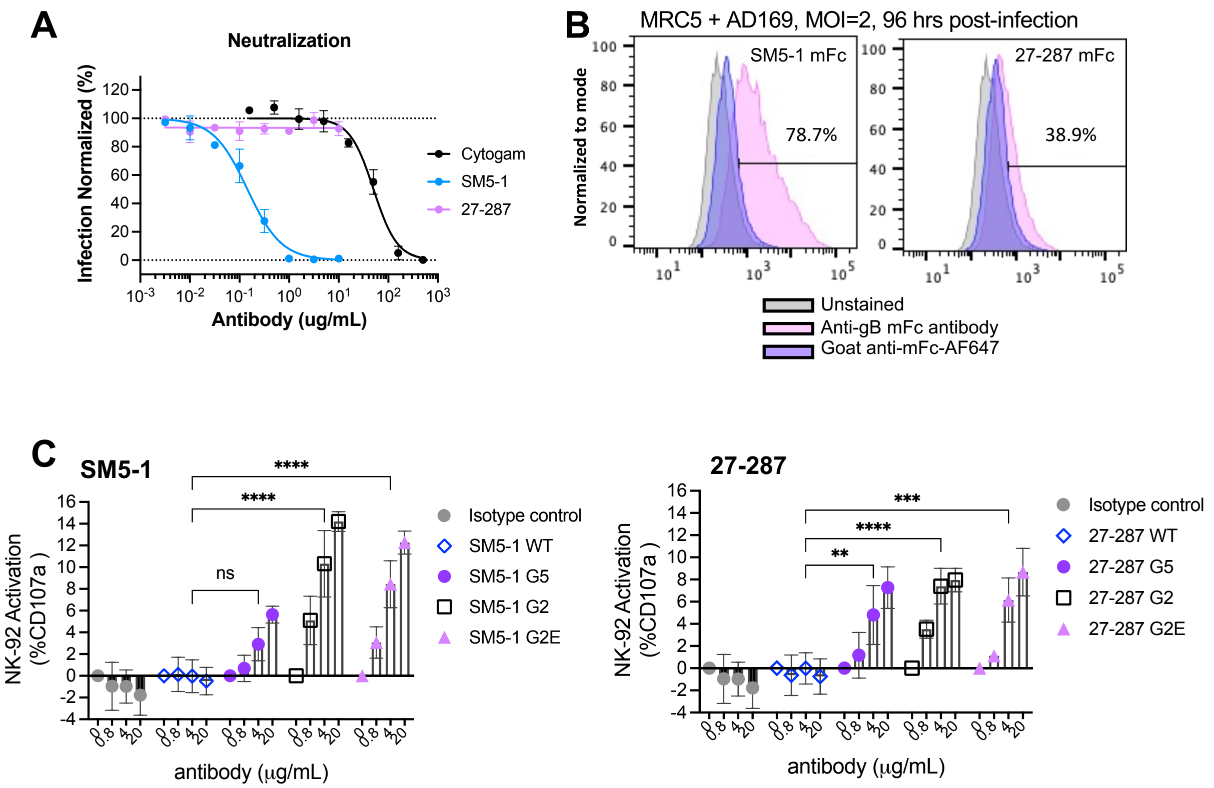


**Figure S12. Anti-gB antibodies 27-287 and SM5-1. A,** Neutralization of AD169 infection of MRC-5 cells by Cytogam, SM5-1 and 27-287, all with WT Fc. Data were normalized to no antibody (100%) and uninfected (0%) controls within each plate and are presented as mean ± SD (n=2) for n=2 experiments. **B,** AD169-MRC5 infected cells at MOI= 2 and 96 hpi were stained with 27-287-mFc or SM5-1-mFc antibodies, with antibody binding measured by flow cytometry. **C,** The percent of NK-92 cells (CD16A V158) that exhibited CD107A-positive degranulation in the presence of AD169-infected MRC5 (MOI=2, 96 hpi) and 27-287 antibodies with WT, G2 and G2E are shown. Data are presented as mean ± SD (n=2) and average of n=3 experiments, two-way ANOVA with Tukey's multiple comparisons test. *$p<0.05$, **$p<0.01$, ***$p<0.001$, ****$p<0.0001$, ns = non-significant.

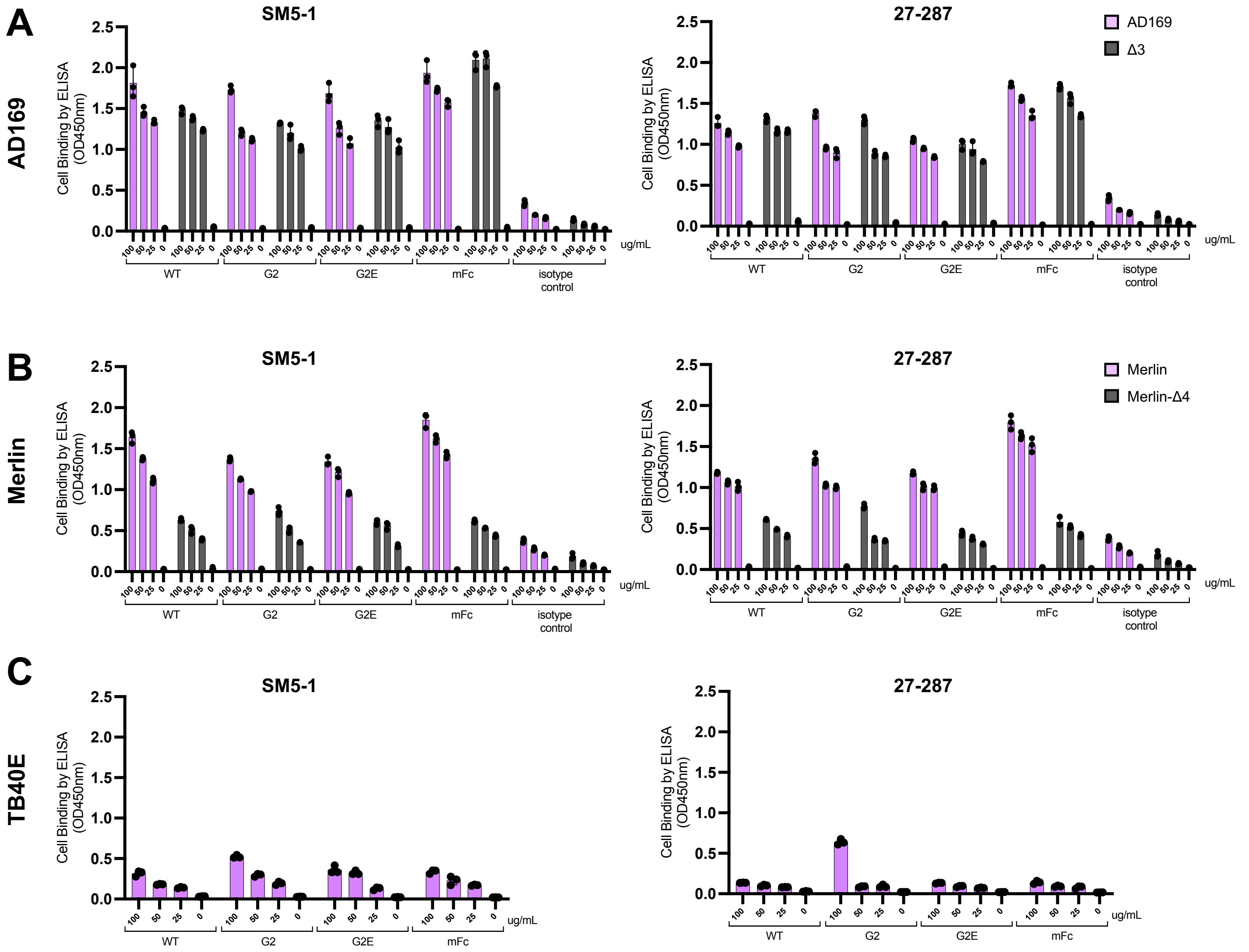


**Figure S13. Comparison of gB binding to HCMV-infected cells**. HFF fibroblast cells were infected with **A,** AD169 or Δ3 and **B,** Merlin or Merlin-Δ4 and **C,** TB40E each at an MOI = 3, followed by CD16A reporter cell assay performed as described in Fig. 5 at 96 hpi. Supernatants removed transferred to another plate for use in mIL2 ELISA measurement and the remaining cells used to measure bound antibodies by goat-anti human IgG-HRP or goat anti-mouse IgG-HRP, respectively by ELISA. Data shown are for three technical replicates from one representative experiment.

**Table S1. Oligonucleotides used in Fc yeast display, generation of AD169 mutants, and viral Fc receptor isolation for CHO cell display.**

| Name | 5’ to 3’ Oligonucleotide | Description |
|---|---|---|
| **Fc cloning into yeast** | | |
| 5’ hinge $CH_2$ | GGTGGTTCTGCTAGCGACAAAACTCAC | Forward primer for Fc IgG1 cloning into pCTcon2 |
| 3’ $CH_3$ | ACTGTTGTTATCAGATCTCGAGCTATTACAAGTCCTCTTCAGAAATAAGCTTTTGTTCGGATCCctatttacccggagacagggagaggctct | Reverse primer for Fc IgG1 cloning into pCTcon2 |
| 3’ $CH_2$ | GGTGTACACCTGTGGTTCTCGGGGCTGCCC | Reverse primer for $CH_2$ amplification |
| 5’ hinge $CH_2$ long | TACGACGTTCCAGACTACGCTCTGCAGGCTAGTGGTGGTGGTGGTTCTGGTGGTGGTGGTTCTGGTGGTGGTGGTTCTGCTAGCgacaaaactcacacatgcccaccgtgcccagcacct | Forward primer for Fc IgG1 cloning into pCTcon2 |
| 3’ $CH_2$ long | ctttgaccaggcaggtcaggctgacctggttcttggtcagctcatcccgggatgggggcagggtgtacacctgtggttctcggggctgccc | Reverse primer for Aga2-CH2 |
| **AD169 mutants** | - | |
| KL-DeltaTRL11 Kana1 | ACGACGAAGAGGACGAGGACGACAACGTCTGATAAGGAAGGCGAGAACGTGTTTTGCACCCCAGTGAATTCGAGCTCGGTAC | Forward primer deletion of *RL11* |
| KL-DeltaTRL11-Kana2 | TGTATACGCCGTATGCCTGTACGTGAGATGGTGAGGTCTTCGGCAGGCGACACGCATCTTGACCATGATTACGCCAAGCTCC | Reverse primer deletion of *RL11* |
| KL-DeltaTRL12-Kana1 | CGGACGGACCTAGATACGGAACCTTTGTTGTTGACGGTGGACGGGGATTTACAGTAAAAGCCAGTGAATTCGAGCTCGGTAC | Forward primer deletion of *RL12* |
| KL-DeltaTRL12-Kana1 | CCTTACAGAATGTTTTAGTTTATTGTTCAGCTTCATAAGATGTCTGCCCGGAAACGTAGCGACCATGATTACGCCAAGCTCC | Reverse primer deletion of *RL12* |
| KL-DeltaUL119-Kana1 | TTGTTTATTTTGTTGGCAGGTTGGCGGGGGAGGAAAAGGGGTTGAACAGAAAGGTAGGTGCCAGTGAATTCGAGCTCGGTAC | Forward primer deletion of *UL118-119* |
| KL-DeltaUL119-Kana2 | AGGTGACGCGACCTCCTGCCACATATAGCTCGTCCACACGCCGTCTCGTCACACGGCAACGACCATGATTACGCCAAGCTCC | Reverse primer deletion of *UL118-119* |
| **vFcγR ectodomains** | - | |
| 5' AD169 *RL12* | ctgcaaccggtgtacactcgAATAGCACCACAACGA | Forward *RL12*, amplify for display on CHO, c-term flag tag |
| 3' AD169 *RL12* | TCCGCCGCTAGCTGAACCACCTCCCTTATCGTCGTCATCCTTGTAGTCAGATCCACGCGGCAGCTCGAGCCGAGAGCGCTGGCTTGAATG | Reverse *RL12*, amplify for display on CHO, c-term flag tag |
| 5’ AD169 *RL11* | caactgcaaccggtgtacactcgAGTTCATCGAACGCCGTCGAA | Forward *RL11*, amplify for display on CHO, c-term flag tag |
| 3’ AD169 *RL11* | GATCCGCCGCTAGCTGAACCACCTCCCTTATCGTCGTCATCCTTGTAGTCAGATCCACGCGGCAGCTCGAGTGAGAGGCCGGACCACTGGCGTTTT | Reverse *RL11*, amplify for display on CHO, c-term flag tag |
| 5’ AD169 *UL118-119* | actgcaaccggtgtacactcgTCAAGTACAACGAGT | Forward *UL118-119*, amplify for display on CHO, c-term flag tag |
| 3’ AD169 *UL118-119* | CGATCCGCCGCTAGCTGAACCACCTCCCTTATCGTCGTCATCCTTGTAGTCAGATCCACGCGGCAGCTCGAGAAGGCGATCCTCGAACAACGGGT | Reverse *UL118-119*, amplify for display on CHO, c-term flag tag |
| 5’ Merlin *RL13* | aactgcaaccggtgtacactcgTATAATCAGACGTGTCC | Forward *RL13*, amplify for display on CHO, c-term flag tag |
| 3’ Merlin *RL13* | ATCCGCCGCTAGCTGAACCACCTCCCTTATCGTCGTCATCCTTGTAGTCAGATCCACGCGGCAGCTCGAGGTTCGCTTTTTTAATGGTTT | Reverse *RL13*, amplify for display on CHO, c-term flag tag |

**Table S2. Pharmacokinetics and thermostabilities of Fc variants**

| | Fc residue changes | % aggregate after thermal stress | $t_{1/2}$ (days) | Clearance (mL/ day) |
|---|---|---|---|---|
| **WT** | N/A | 2.0 ± 0.8 | 8.8 ± 0.9 | 0.30 ± 0.11 |
| **YTE** | M252Y, S254T, T256E | N/A | 12.0 ± 2.6 * | 0.24 ± 0.09 |
| **IHH** | I253A/H310A/H435A | N/A | 0.22 ± 0.01 **** | 4.30 ± 0.35 **** |
| **G5** | S337F, R255Q | ND | 5.1 ± 1.4 * | 0.43 ± 0.08 |
| **G2** | H268L, E294K, Q311L, K334E, Y407V + R255Q | 90.7 ± 0.9*** | 2.2 ± 0.2 **** | 1.26 ± 0.37 **** |
| **G2E** | H268L, R255Q, Q311L, S337F, and K334E | 36.0 ± 2.0*** | 7.6 ± 0.7 | 0.32 ± 0.14 |

Data shown are mean ± SD and compared to WT; statistical significance determined from a one-way ANOVA performed with GraphPad: **$p < 0.01$, ***$p < 0.001$, ****$p < 0.0001$. N/A: not-available. For thermal stress, 1 mg/ml protein was incubated for 20 hrs at 50$^{o}$C.
